# Physical basis for distinct basal and mechanically-gated activity of the human K^+^ channel TRAAK

**DOI:** 10.1101/2021.06.04.447000

**Authors:** Robert A. Rietmeijer, Ben Sorum, Baobin Li, Stephen G. Brohawn

## Abstract

TRAAK is a mechanosensitive two-pore domain K^+^ (K2P) channel localized to nodes of Ranvier in myelinated neurons. TRAAK deletion in mice results in mechanical and thermal allodynia and gain-of-function mutations cause the human neurodevelopmental disorder FHEIG. TRAAK displays basal and stimulus-gated activities typical of K2Ps, but the mechanistic and structural differences between these modes are unknown. Here, we demonstrate that basal and mechanically-gated openings are distinguished by their conductance, kinetics, and structure. Basal openings are low conductance, short duration, and occur through a channel with an interior cavity exposed to the surrounding membrane. Mechanically-gated openings are high conductance, long duration, and occur through a channel that is sealed to the surrounding membrane. Our results explain how dual modes of activity are produced by a single ion channel and provide a basis for the development of state-selective pharmacology with the potential to treat disease.

## Introduction

Two-pore domain K^+^ (K2P) channels display low basal activity (also referred to as background, leak-type, flux-gated, or resting activity) that is important for setting and maintaining the resting membrane potential in many cells (Enyedi and Czirják, 2010; Renigunta et al., 2015). In addition, K2Ps are further activated by diverse physical and chemical stimuli including membrane tension, temperature, voltage, pH, signaling lipids, and calcium to regulate cellular electrical excitability. K2Ps are physiologically relevant targets of volatile anesthetics and antidepressants and their dysregulation has been implicated in pathologies including migraine, depression, pulmonary hypertension, atrial fibrillation, diabetes, Birk-Barel syndrome, and FHEIG syndrome (Facial Dysmorphism, Hypertrichosis, Epilepsy, Intellectual disability, and Gingival outgrowth) (Enyedi and Czirják, 2010;,Barel et al., 2008; Bauer et al., 2018; Heurteaux et al., 2006; Lafrenière et al., 2010; Ma et al., 2013; Royal et al., 2019; Schmidt et al., 2015; Vierra et al., 2015)

TRAAK is a mechanosensitive K2P expressed in the nervous system of jawed vertebrates and localized to nodes of Ranvier, the gaps in myelinated axons where the action potential is regenerated during saltatory conduction. TRAAK accounts for ∼25% of nodal background K^+^ conductance and maintains the resting membrane potential and voltage gated-Na^+^ channel availability required for high frequency spiking (Brohawn et al., 2019; Kanda et al., 2019;,Maingret et al., 1999). Altering TRAAK activity in animals has physiological consequences; TRAAK knockout mice display mechanical and thermal allodynia and mechanical hyperalgesia while gain-of-function mutations in humans (TRAAK_A198E_ and TRAAK_A270P_) underlie the neurodevelopmental disorder FHEIG (Bauer et al., 2018; Noël et al., 2009).

TRAAK basal activity can be activated ∼100-fold by increased membrane tension (Brohawn et al., 2014a, 2014b; Sorum et al., 2021). A model for mechanical gating primarily involves conformational changes in transmembrane helix 4 (TM4) (Brohawn et al., 2014b). In this model, nonconductive (closed) TRAAK adopts a TM4 “down” conformation and conductive (open) TRAAK adopts a TM4 “up” conformation. When TM4 is down, a lipid acyl chain can enter the channel cavity below the selectivity filter through membrane facing openings and sterically block the conduction path. Movement of TM4 up seals the membrane facing openings, prevents lipid access, and permits ion conduction. Structures of these two conformations provide a biophysical explanation for mechanosensitivity: shape changes upon opening, including an expansion in cross sectional area and increase in cylindricity, are energetically favored by membrane tension.

Notably, single channel recordings of TRAAK, and the closely related mechanosensitive K2P channels TREK1 and TREK2, suggest the presence of multiple distinct open and closed states (Bang et al., 2000; Kang et al., 2005; Maingret et al., 1999; Patel et al., 1998; Sorum et al., 2021; Xian Tao Li et al., 2006; Clausen et al. 2020). In addition, alternative models for gating mechanosensitive K2Ps posit closures involving C-type gating at the selectivity filter (Chatelain et al., 2012; González et al., 2013; Li et al., 2020; Lolicato et al., 2020; Schewe et al., 2016), closures involving dewetting of the channel cavity (Aryal et al., 2014, 2015), and openings that do not involve upward movement of TM4 (Lolicato et al., 2014). For TRAAK, and other K2Ps, basally open and stimulus-activated currents display different voltage dependence (Schewe et al., 2016), consistent with structurally distinct open states. Here, we show basal and mechanically-gated TRAAK open states are distinct in their conductance, kinetics, and structure and present a model for channel gating that associates conformational and functional states.

## Results and Discussion

### Basal and mechanically-gated TRAAK openings are physically distinct

A macroscopic recording from a patch containing wild-type TRAAK (TRAAK_WT_) illustrates basal activity and robust activation by membrane tension (generated by negative pressure applied through the patch pipette) that are characteristic of the channel (Fig. 1A,B). We performed single channel recordings to determine whether basally open and mechanically activated TRAAK have different biophysical characteristics consistent with distinct channel open states. By varying applied pressure over approximately an hour of single channel records, we sampled a wide range of TRAAK_WT_ activity (0.01 ≤ open probability (P_O_) ≤ 0.98) (Fig. 1C-F, Table S1). We analyzed records in three bins according to P_O_ (Fig. 1C). We found that in addition to a simple increase in P_O_, mechanically activated TRAAK openings are distinct from basal openings in two ways.

**Figure 1.**
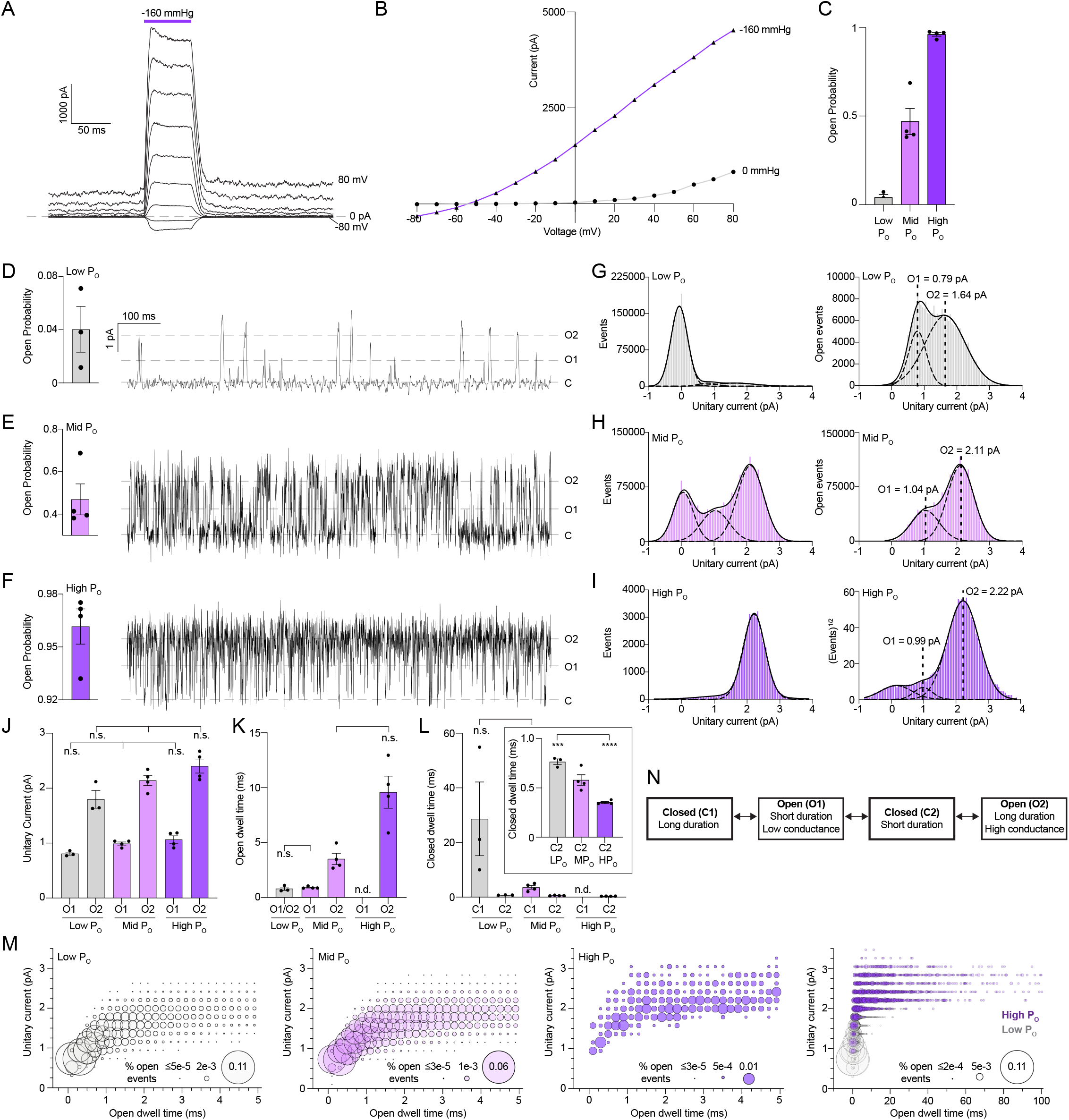
Distinct basal and mechanically-activated open states in TRAAK_WT_. (A) Macroscopic currents recorded across an inside-out patch pulled from a TRAAK-expressing cell in response to a voltage step protocol (V_hold_ = 0, V_test_ = −80 to 80, ΔV = 10 mV, 20 mV intervals displayed) with a pressure step applied at each voltage (purple bar). (B) Current-voltage relationship from (A). (C) Open probability calculated from single channel records (0.04 ± 0.02, 0.47 ± 0.07, and 0.96 ± 0.01 for low P_O_, mid P_O_, and high P_O_, respectively (mean ± sem, n = 3, 4, and 4 patches)). All single channel data in the paper were recorded at V_hold_ = 0 mV in a ten-fold gradient of [K^+^] and are presented in physiological convention. (D-F) 1 s portion from representative low P_O_, mid P_O_, and high P_O_ recordings, respectively. (G-I) All event (left) and open-only or square root all event (right) current histograms from representative low P_O_, mid P_O_, and high P_O_ recordings, respectively. (J) Unitary currents of O1 and O2 states (0.81 ± 0.03 pA and 1.80 ± 0.16 pA, 0.99 ± 0.03 pA and 2.14 ± 0.09 pA, and 1.07 ± 0.07 pA and 2.41 ± 0.13 pA for low P_O_, mid P_O_, and high P_O_ recordings, respectively (mean ± sem, n= 3, 4, and 4 patches)). (K) Open dwell time of O1 and O2 states (0.80 ± 0.15 ms, 0.91 ± 0.05 ms and 3.52 ± 0.51 ms, and 9.60 ± 1.46 ms for low P_O_, mid P_O_, and high P_O_ recordings, respectively (mean ± sem, n= 3, 4, and 4 patches)). (L) Closed dwell times of C1 and C2 states (28.70 ± 13.50 and 0.77 ± 0.03 ms, 3.59 ± 0.67 and 0.58 ± 0.05 ms, and 0.35 ± 0.01 ms for low P_O_, mid P_O_, and high P_O_ recordings, respectively (mean ± sem, n=3, 4, and 4 patches, n.d. not determined, n.s. not significant, ***p = 0.0006, ****p < 0.0001 (one-way ANOVA with Tukey correction)). (M) Unitary current-open dwell time relationships for low P_O_, mid P_O_, and high P_O_ open events and an overlay of low P_O_ and high P_O_ relationships at expanded time scale. Bubble size is proportional to percentage of open events. (N) Four state TRAAK gating model.

First, mechanical force favors higher conductance openings (Figs. 1G-I, S1). Two conductance states are observed in TRAAK_WT_: a low conductance opening with a unitary current of ∼1 pA and a high conductance opening with a unitary current of ∼2 pA. In unstretched patches at low P_O_ (0.01 ≤ P_O_ ≤ 0.07), the two conductance states are observed at roughly similar frequency (Fig. 1G,M). At intermediate P_O_ (mid P_O_, 0.38 ≤ P_O_ ≤ 0.69), more high conductance openings are observed (Fig. 1H,M). In records mechanically activated to high P_O_ (0.93 ≤ P_O_ ≤ 0.98), almost all openings are high conductance (Fig. 1I,M). Notably, unitary currents of these openings do not change significantly as a function of P_O_, but the relative proportion of each conductance state does (Fig. 1G-J,M).

Second, mechanical force favors longer duration openings. Two kinetically distinct open states are observed in TRAAK_WT_ records: a short duration (∼1 ms) opening and a long duration (∼3 ms or longer) opening (Figs. 1K, S2). In unstretched patches at low P_O_, almost all openings are short duration (Fig. 1M). In contrast, when mechanically activated to high P_O_, almost all openings are long duration. At intermediate P_O_, both short and long duration openings are observed. Open duration and unitary current are positively correlated: low conductance openings have a narrow duration distribution around 1 ms while high conductance openings have a wider duration distribution ≥2 ms, with longer openings observed at higher P_O_ (Fig. 1M).

In addition, mechanical force favors shorter duration closures. TRAAK_WT_ accesses two kinetically distinct closed states: a short duration (∼0.5 ms) closure and a long duration (∼3 ms or longer) closure (Figs. 1L, S3). Short duration closures of comparable length are observed in all records. However, long duration closures are only observed in low and intermediate P_O_ records and their prevalence and duration decrease as P_O_ increases (Figs. 1L, S3).

Single channel TRAAK_WT_ records can be fit with a linear kinetic model relating the two closed states (long duration (C1) and short duration (C2)) and two open states (short duration/low conductance (O1) and long duration/high conductance (O2)) (Fig. 1N, Table S1)(Sorum et al., 2021). The distribution of states and the duration of C1 and O2 are dependent on membrane tension. When mechanically activated to high P_O_, long duration closures and short duration/low conductance openings are almost never observed, so data can be fit with a two-state equilibrium model between C2 and O2 states (Figs. 1I,K-M, S1-3, Table S1). At intermediate P_O_, the duration of long C1 closures is shorter than those at low P_O_ and the duration of O2 openings is shorter than those at high P_O_ (Figs. 1H, K-M, S1-3, Table S1). We conclude that mechanically activated openings correspond to the O2 state while basal openings correspond to the O1 state. We speculate weak mechanical activation by resting membrane tension in the patch results in low P_O_ O2 openings with a conductance distinct from O1 openings (Brohawn et al., 2014b;,Opsahl and Webb, 1994). However, the duration of low P_O_ O1 and O2 openings is similarly short and therefore indistinguishable kinetically (Fig. 1D,K).

We then asked whether we could identify the structural basis for these two distinct TRAAK open states. We reasoned that gain-of-function mutations could differentially stabilize particular open states and investigated three point mutants reported to dramatically increase channel activity: TRAAK_A198E_, TRAAK_A270P_, and TRAAK_G158D_. TRAAK_A198E_ and TRAAK_A270P_ were recently identified in human patients with the neurological and developmental disorder FHEIG while TRAAK_G158D_ was identified as a pan-K2P gain-of-function mutation (Bauer et al., 2018; Ben Soussia et al., 2019).

### Mechanical force and FHEIG mutations promote a TM4 up open state

We first consider the FHEIG mutations TRAAK_A198E_ and TRAAK_A270P_ (Figs. 2A, 3A). Macroscopic recordings of TRAAK_A198E_ or TRAAK_A270P_ channels display currents that are only subtly activated by mechanical force, consistent with the initial characterization of these mutations (Figs. 2B,C, 3B,C) (Bauer et al., 2018). Single channel records reveal a simple explanation for reduced mechanical activation of TRAAK_A198E_ and TRAAK_A270P_ relative to TRAAK_WT_. TRAAK_A198E_ and TRAAK_A270P_ open probability is nearly one at rest and mechanical force does not change channel conductance, so stimulation can only slightly increase channel activity (Figs. 2D-G, 3D-G).

**Figure 2.**
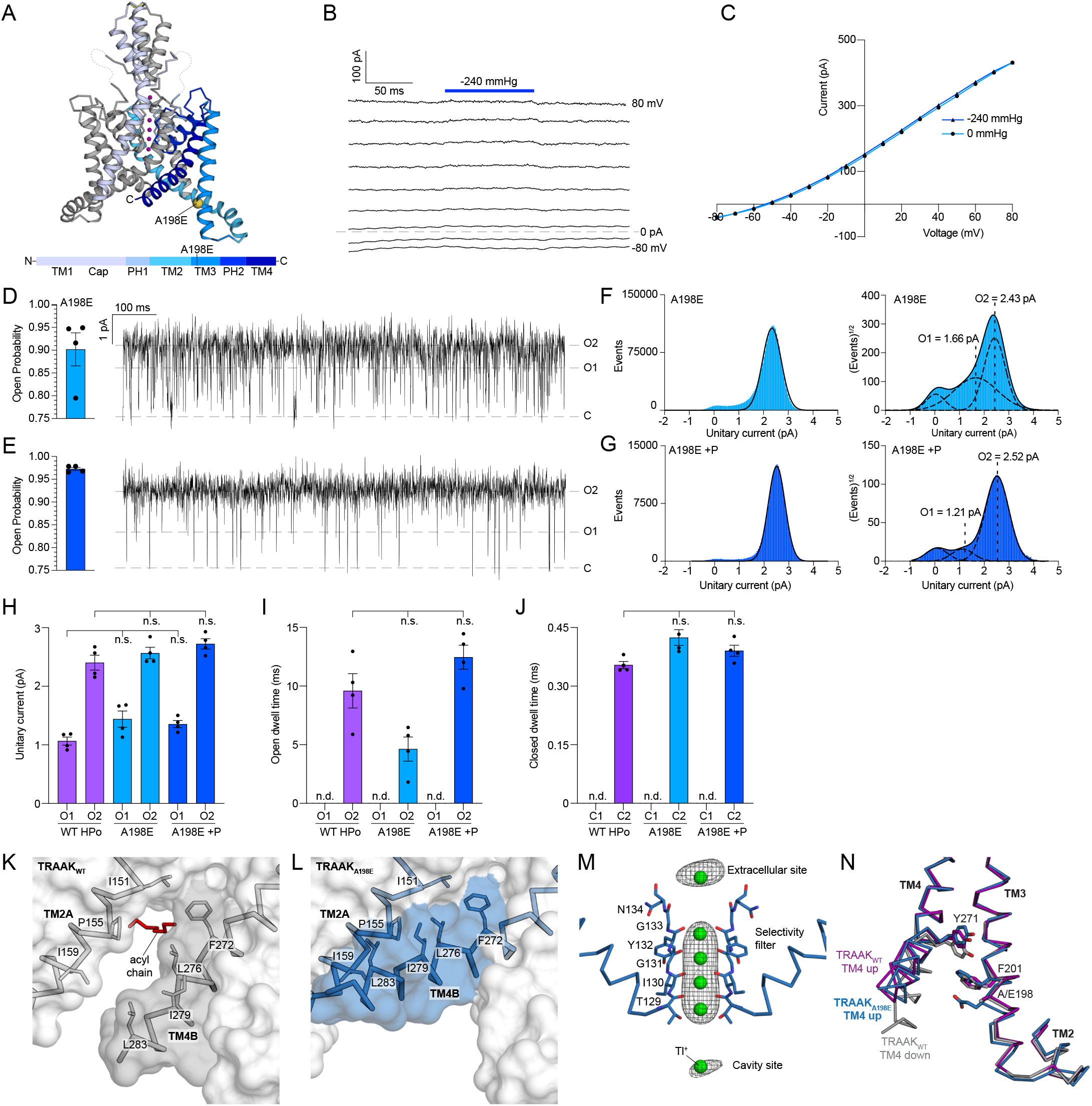
The FHEIG mutation TRAAK_A198E_ promotes a long duration, high conductance TM4-up open state. (A) Crystal structure of TRAAK_A198E_. Side view from the membrane plane with one protomer gray and the second protomer colored according to the key below. A198E is shown as a yellow sphere and K^+^ ions are colored purple. (B) Macroscopic currents from a TRAAK_A198E_-containing patch in response to a voltage step protocol (V_hold_ = 0, V_test_ = −80 to 80, ΔV = 10 mV, 20 mV intervals displayed) with a pressure step applied at each voltage (dark blue bar). (C) Current-voltage relationship from (B). (D,E) Open probability calculated from all TRAAK_A198E_ and TRAAK_A198E_ with pressure (+P) records (left, P_O_ = 0.90 ± 0.04 and P_O_ = 0.97 ± 0.003 (mean ± sem, n = 4 patches) and 1 s portion from representative recordings. (F,G) All event (left) and square root all event (right) current histograms from representative recordings. (H) Unitary currents of TRAAK_WT_ HP_O_, TRAAK_A198E_, and TRAAK_A198E_+P O1 and O2 states (1.07 ± 0.07 and 2.41 ± 0.13 pA, 1.44 ± 0.14 and 2.57 ± 0.10 pA, and 1.35 ± 0.06 and 2.72 ± 0.09 pA, respectively). (I) Open dwell times of TRAAK_WT_HPO, TRAAK_A198E_, and TRAAK_A198E_+P O2 states (9.60 ± 1.46 ms, 4.63 ±1.03 ms, 12.46 ±1.02 ms, respectively). (J) Closed dwell times of TRAAK_WT_ HPO, TRAAK_A198E_, and TRAAK_A198E_+P C2 (0.35 ± 0.01 ms, 0.42 ± 0.02 ms, and 0.39 ± 0.01 ms, respectively). For H-J, data are mean ± sem, n = 4 patches; n.d., not determined; n.s., not significant (one-way Anova with Dunnett correction). (K) View of the membrane-facing lateral opening in a TRAAK_WT_ TM4-down structure (PDB 4WFF). A cavity-bound lipid acyl chain blocks conduction. (L) TRAAK in the same view as (K). A TM4-up conformation seals the membrane opening. (M) Ions in a TRAAK_A198E_ -Tl^+^ structure. Anomalous density (grey) around Tl+ ions (green) displayed at 2.5 σ (extracellular / selectivity filter / cavity ions). (N) Overlay of the TM2-TM3-TM4 interaction from TRAAK_A198E_ and TRAAK_WT_ structures. A198E sterically promotes a TM4-up open state.

TRAAK_A198E_ and TRAAK_A270P_ single channel behavior closely matches TRAAK_WT_ when mechanically activated to high P_O_. TRAAK_A198E_ and TRAAK_A270P_ openings are nearly all high conductance (with a unitary current ∼2 pA) and long duration (∼3 ms or longer), while closures are short duration (∼0.5 ms) (Figs. 2F-J, 3F-J). The modest mechanical activation of TRAAK_A198E_ and TRAAK_A270P_ is due to an increase in long duration openings without significantly changing closed duration.

We determined crystal structures of TRAAK_A198E_ and TRAAK_A270P_ in the presence of K^+^ and in complex with a mouse monoclonal antibody Fab fragment to 2.5 Å and 3.0 Å resolution, respectively (Figs. 2A, 3A, S4). As previously reported for this crystal form, the conformation of one side of the TRAAK dimer is restricted because it is involved in forming contacts that propagate the lattice (Fig. S5). We therefore focus our discussion on the opposing side of the TRAAK dimer (including TM4 from protomer B), which is conformationally unrestricted. In the same crystallization conditions, TRAAK_WT_ adopts a TM4 down nonconductive conformation; the membrane-facing lateral opening above TM4 permits hydrophobic acyl chains to access the channel cavity and block ion passage (Fig. 2K) (Brohawn et al., 2014b). TRAAK_A198E_ and TRAAK_A270P_, however, adopt TM4 up conformations (Figs. 2L, 3K). Movement of TM4 up seals the membrane-facing lateral openings, preventing acyl chain access to the cavity and creating an unobstructed path for ion movement through the channel. This TM4 up conformation is similar to a TRAAK_WT_ TM4 up conductive conformation captured in the presence of a small molecule activator trichloroethanol (Fig. S7B,E) (Brohawn et al., 2014b). As observed in the TRAAK_WT_ TM4 up conductive structure, density consistent with a K^+^ ion is present in the cavities of TRAAK_A198E_ and TRAAK_A270P_ (Figs. 3L, S6A).

**Figure 3.**
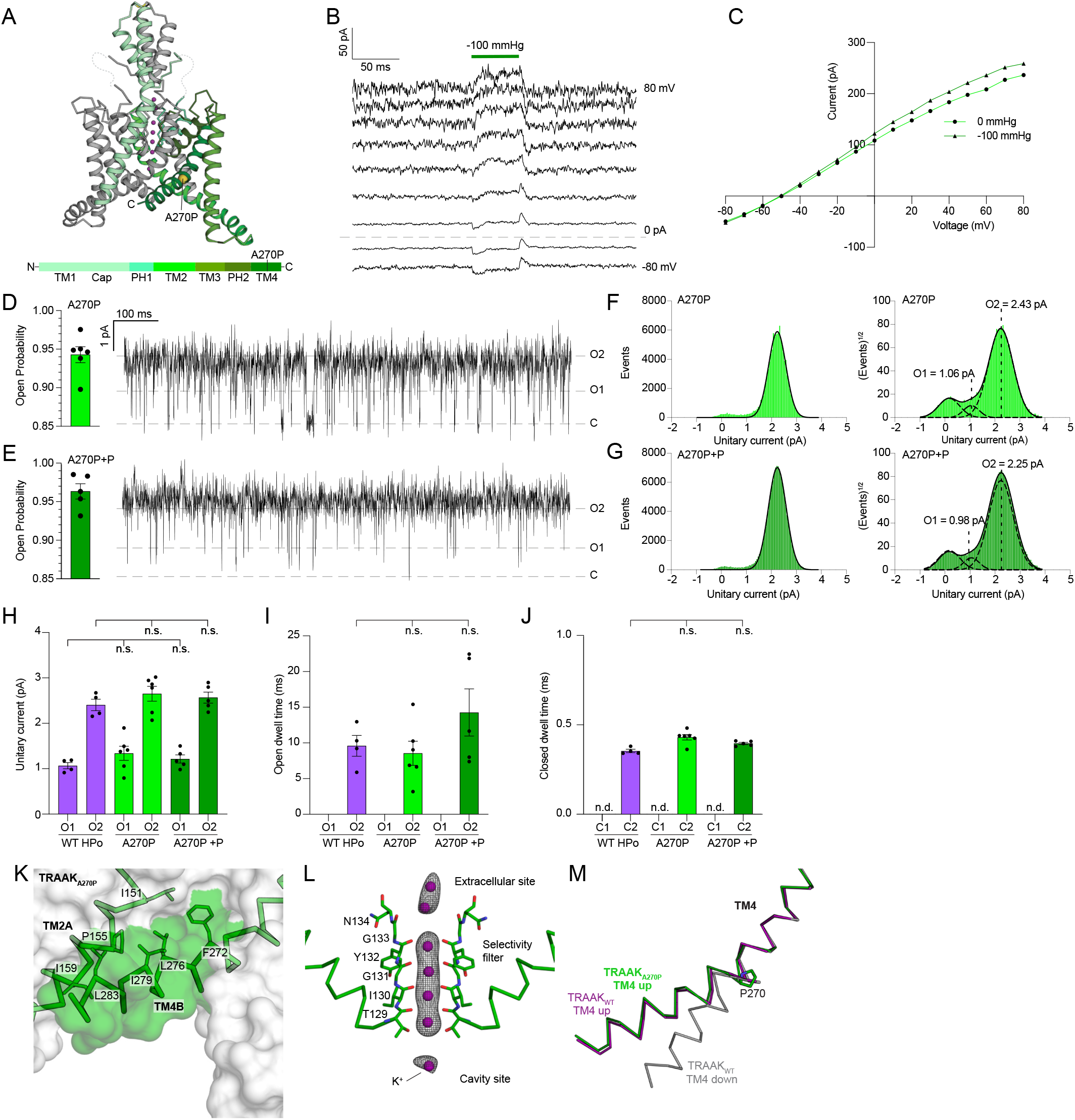
The FHEIG mutation TRAAK_A270P_ promotes a long duration, high conductance TM4-up open state. (A) Crystal structure of TRAAK_A270P_. Side view from the membrane plane with one protomer gray and the second protomer colored according to the key below. A270P is shown as a yellow sphere and K^+^ ions are colored purple. (B) Macroscopic currents from a TRAAK_A270P_-containing patch in response to a voltage step protocol (V_hold_ = 0, V_test_ = −80 to 80, ΔV = 10 mV, 20 mV intervals displayed) with a pressure step applied at each voltage (dark green bar). (C) Current-voltage relationship from (B). (D,E) Open probability calculated from all TRAAK_A270P_ and TRAAK_A270P_ with pressure (+P) records (left, P_O_ = 0.94 ± 0.01 and P_O_ = 0.96 ± 0.01 (mean ± sem, n = 6 and 5 patches) and 1 s portion from representative recordings. (F,G) All event (left) and square root all event (right) current histograms from representative recordings. (H) Unitary currents of TRAAK_WT_ HP_O_, TRAAK_A270P_, and TRAAK_A270P_+P O1 and O2 states (1.07 ± 0.07 and 2.41 ± 0.13 pA, 1.34 ± 0.15 and 2.65 ± 0.16 pA, and 1.21 ± 0.09 and 2.57 ± 0.12 pA, respectively). (I) Open dwell times of TRAAK_WT_HP_O_, TRAAK_A270P_, and TRAAK_A270P_+P O2 states (9.60 ± 1.46 ms, 8.56 ± 1.69 ms, and 14.47 ± 3.30 ms, respectively). (J) Closed dwell times of TRAAK_WT_ HP_O_, TRAAK_A270P_, and TRAAK_A270P_+P C2 (0.35 ± 0.01 ms, 0.43 ± 0.01 ms, and 0.39 ± 0.01 ms, respectively). For H-J, data are mean ± sem, n = 4, 6, and 5 patches; n.d., not determined; n.s., not significant (one-way Anova with Dunnett correction). (K) View of the membrane-facing cytoplasmic half of TM4 in TRAAK_A270P_. A TM4-up conformation seals the membrane opening. (L) Ions in the TRAAK _A270P_ -K^+^ structure. Polder omit F_o_-F_c_ density (grey) around K^+^ ions (purple) displayed at 5 and 5.5 σ for extracellular and selectivity filter and cavity ions, respectively. (M) Overlay of TM4 from TRAAK_A270P_ and TRAAK_WT_ structures. A270P kinks TM4 to promote a TM4-up open state.

To confirm that the TRAAK_A198E_ TM4 up structure represents an open state, we determined a second structure in the presence of Tl^+^ to 3.0 Å resolution and used anomalous diffraction of Tl^+^ ions to unambiguously identify ion binding sites. Indeed, a Tl^+^ ion is identified in the channel cavity in addition to four sites in the selectivity filter (S1-S4) and one on the extracellular mouth of the pore (S0), consistent with a conductive conformation (Fig. 2M).

How do TRAAK_A198E_ and TRAAK_A270P_ promote TM4 up open states? Comparing TRAAK_A198E_ and TRAAK_WT_ suggests two mechanisms in this mutant (Figs. 2N, S6D&E). The first is a steric relay from A198E through F201 and Y271 that favors TM4 up. In TRAAK_WT_, the TM4 down conformation involves movement of Y271 1 Å towards the cytoplasm relative to TM4 up. This requires the intracellular TM2-TM3 linkage to rotate 15° away from TM4 to prevent a clash between Y271 on TM4 and F201 on TM3. In TRAAK_A198E_, TM2-TM3 is similarly rotated out, but A198E pushes F201 closer to TM4 where it would clash with Y271 in a TM4 down conformation. A second possibility is that A198E causes local thinning of the lipid bilayer to favor TM4 up. While A198 is predicted to be embedded in the membrane inner leaflet facing hydrophobic residues on TM4 down, the negatively charged carboxylic acid of A198E likely favors hydration of this pocket, promoting TM4 up to maintain nonpolar interactions between its hydrophobic residues and lipid acyl chains (Brooks, B. R. and Brooks, III, C. L.1 and Mackerell, Jr. et al., 2009; Jo et al., 2008).

In TRAAK_A270P_, the introduced proline kinks TM4 to closely approximate the TRAAK_WT_ TM4 up conformation (Fig 3M). In TRAAK_WT_, movement of TM4 up depends on a hinge at a conserved glycine G268. The A270P mutation instead disrupts backbone hydrogen bonding in TM4 and creates a ∼27° bend and ∼9.5 Å upward movement of TM4 relative to TRAAK_WT_ TM4 down.

We conclude that the TRAAK_A198E_, TRAAK_A270P_, and TRAAK_WT_ TM4 up structures correspond to the high conductance/long duration mechanically activated open state O2. The following lines of evidence support this assignment: (i) TRAAK_A198E_ and TRAAK_A270P_ have an open probability ≥ 0.9 (Figs. 2F,H, 3F,H), (ii) the structures are similar to a TRAAK_WT_ mechanically activated conductive conformation (Figs. 2K,L, 3K), (iii) the structures show unobstructed conduction paths with ions in the channel cavities (Figs. 2M, 3L, S6A), (iv) the predominant TRAAK_A198E_ and TRAAK_A270P_ unitary current is indistinguishable from TRAAK_WT_ O2 (Figs. 1I, 2G, 3G), and (v) the mean duration of the TRAAK_A198E_ and TRAAK_A270P_ open states are indistinguishable from TRAAK_WT_ O2 (Figs. 1K, 2I, 3I, S2).

### A TM4 down open state promoted by a pan-K2P activating mutation underlies basal activity

We next consider TRAAK_G158D_ (Fig. 4A). In macroscopic recordings, mechanical force activates TRAAK_G158D_ ∼2-fold; more than TRAAK_A198E_ and TRAAK_A270P_, but less than TRAAK_WT_ (Fig. 4B-D). Single channel analysis shows that, similar to TRAAK_WT_, mechanical activation of TRAAK_G158D_ involves an increase in both open probability and conductance (Fig. 4E-H). However, the resting open probability of TRAAK_G158D_ (P_O_ ∼0.7) is much higher than TRAAK_WT_ (P_O_ ∼0.04) such that mechanical stimulation can only increase TRAAK_G158D_ P_O_ ∼1.25 fold (Fig. 4E,F). Basal openings of TRAAK_G158D_ are low conductance with a unitary current ∼1 pA while mechanically activated openings are higher conductance with a unitary current ∼2 pA (Figs. 4G-I, S1).

**Figure 4.**
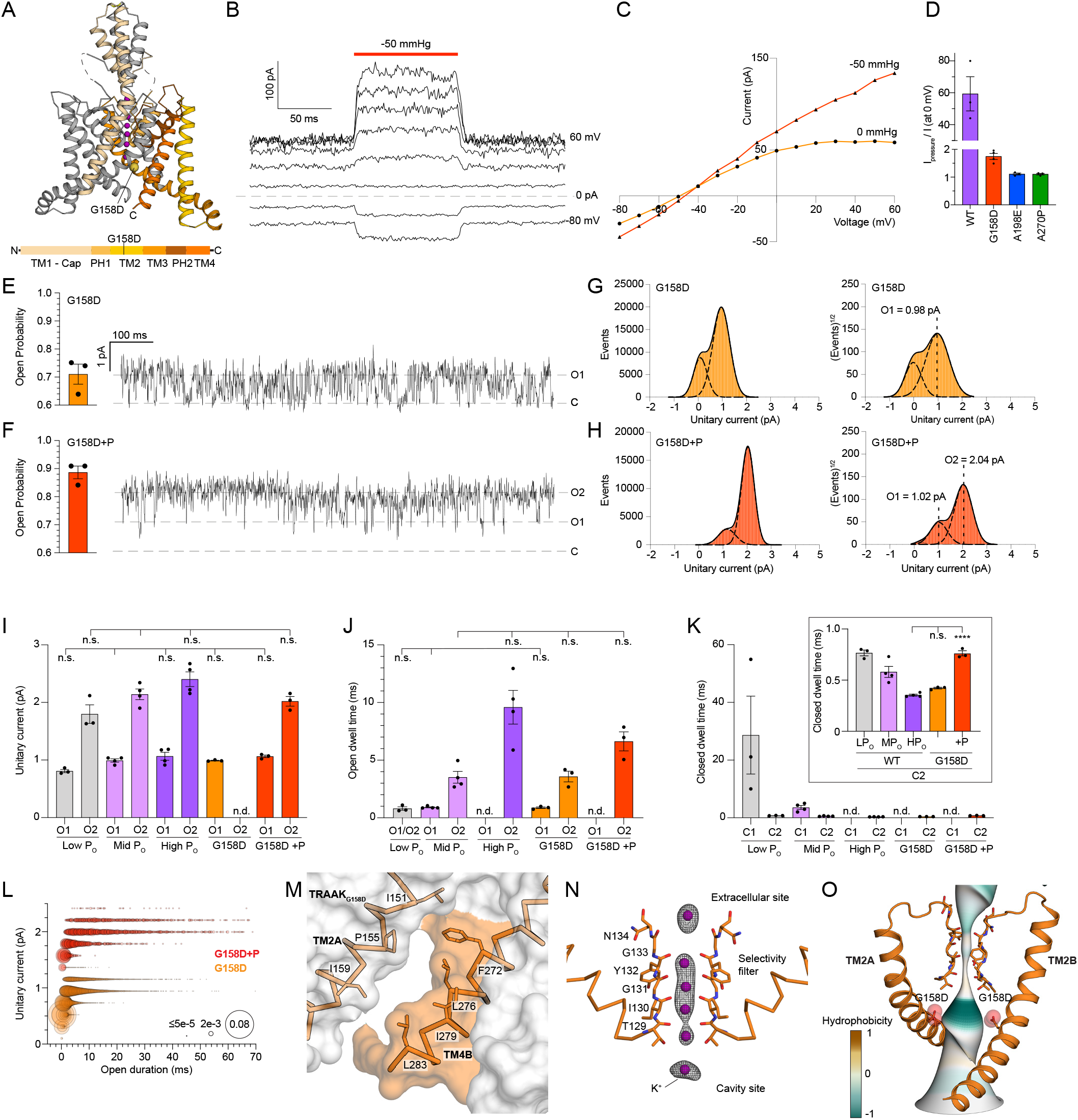
The gain-of-function mutation TRAAK_G158D_ promotes a short duration, low conductance TM4-down open state. (A) Crystal structure of TRAAK_G158D_. Side view from the membrane plane with one protomer gray and the second protomer colored according to the key below. G158D is shown as a yellow sphere and K^+^ ions are colored purple. (B) Macroscopic currents from a TRAAK_G158D_-containing patch in response to a voltage step protocol (V_hold_ = 0, V_test_ = −80 to 80, ΔV = 10 mV, 20 mV intervals displayed) with a pressure step applied at each voltage (dark orange bar). (C) Current-voltage relationship from (B). (D) Maximum fold-activation by pressure of macroscopic TRAAK_WT_, TRAAK_G158D_, TRAAK_A198E_, and TRAAK_A270P_ currents with expanded scale in inset (59.33 ± 10.73, 1.75 ± 0.11, 1.11 ± 0.03, and 1.10 ± 0.03, respectively, mean ± sem, n = 3, 4, 3, and 3). (E,F) Open probability calculated from all TRAAK_G158D_ and TRAAK_G158D_ with pressure (+P) records (left, P_O_ = 0.71 ± 0.04 and P_O_ = 0.89 ± 0.02 (mean ± sem, n=3 patches) and 1s portion from representative recordings. (G,H) All event (left) and square root all event (right) current histograms from representative recordings. (I) Unitary currents of TRAAK_WT_ LP_O_, MP_O_, HP_O_, TRAAK_G158D_, and TRAAK_G158D_+P O1 and O2 states (0.81 ± 0.03 pA and 1.80 ± 0.16 pA, 0.99 ± 0.03 pA and 2.14 ± 0.09 pA, 1.07 ± 0.07 pA and 2.41 ± 0.13 pA, 0.99 ± 0.01, and 1.06 ± 0.02 and 2.02 ± 0.08 pA, respectively). (J) Open dwell times of TRAAK_WT_ LP_O_, MP_O_, HP_O_, TRAAK_G158D_, and TRAAK_G158D_+P O1 and O2 states (0.80 ± 0.15 ms, 0.91 ± 0.05 ms and 3.52 ± 0.51 ms, 9.60 ± 1.46 ms, 0.88 ± 0.07 and 3.58 ± 0.46 ms, and 6.63 ± 0.83 ms, respectively). (K) Closed dwell times of TRAAK_WT_ LP_O_, MP_O_, HP_O_, TRAAK_G158D_, and TRAAK_G158D_+P C1 and C2 states (28.70 ± 13.50 and 0.77 ± 0.03 ms, 3.59 ± 0.67 and 0.58 ± 0.05 ms, 0.35 ± 0.01 ms, 0.42 ± 0.01 ms, and 0.76 ± 0.03 ms, respectively). For I-K, data are mean ± sem, n = 3, 4, 4, 3 and 3 patches; n.d., not determined; n.s., not significant, ****p < 0.0001 (one-way Anova with Tukey correction). (L) Unitary current-open dwell time relationships for TRAAK_G158D_, and TRAAK_G158D_+P open events and an overlay at expanded time scale. Bubble size is proportional to percentage of open events. (M) View of the membrane-facing cytoplasmic half of TM4 in TRAAK_G158D_. (N) Ions in the TRAAK _G158D_ -K^+^ structure. Polder omit F_o_-F_c_ density (grey) around K^+^ ions (purple) displayed at 6.5 and 6 σ for selectivity filter and extracellular and cavity ions, respectively. (O) The conduction path in TRAAK_G158D_ colored by hydrophobicity. G158D increases cavity electronegativity to promote a TM4-down open state.

Kinetic analysis shows that TRAAK_G158D_ closely resembles TRAAK_WT_ except that the long duration closed state C1 is absent in the mutant (Figs. 4J, S2, S3). TRAAK_G158D_ accesses only a single short closed state (Fig. 4K). Like TRAAK_WT_, TRAAK_G158D_ has two open states, one short duration/low conductance (∼1 ms / 1 pA) and one long duration/high conductance (∼3 ms or longer / 2 pA) (Fig. 4I,J). Mechanical stimulation favors longer duration/high conductance openings. (Figs. 4L, S1, S2).

We determined the crystal structure of TRAAK_G158D_ to 3.0 Å resolution in the presence of K^+^ (Fig. 4A). TRAAK_G158D_ adopts a TM4 down conformation like TRAAK_WT_ crystallized under the same conditions (Fig. 4M). However, density in the TRAAK_G158D_ cavity is consistent with a K^+^ ion rather than the hydrophobic acyl chain present in the TRAAK_WT_ TM4 down structure (Fig. 4N). This suggests that the TRAAK_G158D_ structure represents a TM4 down open state and is consistent with the high resting open probability of the mutant channel.

TRAAK_G158D_ likely promotes a TM4 down open state through a simple electrostatic consequence of the mutation (Fig. 4O). G158D projects into the channel cavity ∼5 Å underneath the selectivity filter and increases hydrophilicity of the ion conduction path close to where acyl chains are observed in nonconductive TM4 down TRAAK_WT_ structure. The electronegative cavity of TRAAK_G158D_ is expected to disfavor acyl chain access and channel block, instead promoting ion occupancy in the cavity and a conductive open state.

We conclude that the TM4 down open state captured in the TRAAK_G158D_ structure corresponds to the low conductance/short duration basal open state O1 in TRAAK_WT_. The following data support this assignment: (i) TRAAK_G158D_ has a high resting open probability (Fig. 4E), (ii) a K^+^ ion rather than an acyl chain is observed in the TRAAK_G158D_ cavity (Fig. 4N), (iii) unitary current of the TRAAK_G158D_ basal open state is indistinguishable from TRAAK_WT_ O1 (Fig. 4I), (iv) mean dwell time of the TRAAK_G158D_ short duration opening is statistically indistinguishable from TRAAK_WT_ O1 (Fig. 4J), and (v) like TRAAK_WT_, the basal low conductance/short duration TRAAK_G158D_ O1-like open state can be mechanically activated to a high conductance/long duration O2-like open state (Fig. 4F-J). Consistent with (v), TRAAK_G158D_ can adopt a TM4 up-like conformation (observed on the side of the channel involved in forming crystal contacts) and TRAAK_G158D_ TM4 up is expected to be energetically favored in the presence of membrane tension relative to TRAAK_G158D_ TM4 down because it has an increased cross-sectional area (Fig. S7C). The lower conductance of TRAAK O1 openings may be due to reduced cavity polarity or a longer path length through the channel in a TM4-down conformation relative to the TM4-up conformation of O2 openings.

### An integrated structural and functional model for TRAAK gating

We further conclude that the TM4 down lipid-blocked closed state observed in the TRAAK_WT_ structures corresponds to the long duration closed state C1. Analogous long duration closures are not observed in in TRAAK_G158D_, TRAAK_A198E,_ or TRAAK_A270P_ (Figs. 1L, 2J, 3J, 4K). In TRAAK_A198E_ and TRAAK_A270P_, this is likely because TM4s are favored in an up conformation that seals lateral membrane openings, preventing lipid access to the cavity (Figs. 2L, 3K). In TRAAK_G158D_, this is likely because the mutation increases polarity of the channel cavity to disfavor occupancy of hydrophobic acyl chains (Fig. 4O).

Together, these data support a model for TRAAK gating shown in Figure 5 that maps structural conformations to the linear four-state kinetic model derived from single-channel records (Fig. 1N). Long duration closures (C1) correspond to the TM4 down, lipid blocked conformation captured in TRAAK_WT_ structures(Brohawn et al., 2014b). Low conductance/short duration basal openings (O1) correspond to the TM4 down, conductive conformation captured in the TRAAK_G158D_ structure (Fig. 4). High conductance/long duration mechanically activated openings (O2) correspond to the TM4 up, conductive conformation captured in structures of TRAAK_A198E_, TRAAK_A270P_, and TRAAK_WT_ with activating small molecules (Fig. 2,3). Structures of two other gain-of-function mutations, TRAAK_G124I_ and TRAAK_W262S_, have been previously reported(Lolicato et al., 2014), although their open probability, single channel behavior, and cavity ion occupancy is not known. However, both adopt TM4 down-like conformations and may promote O1 like openings similarly to TRAAK_G158D_.

**Figure 5.**
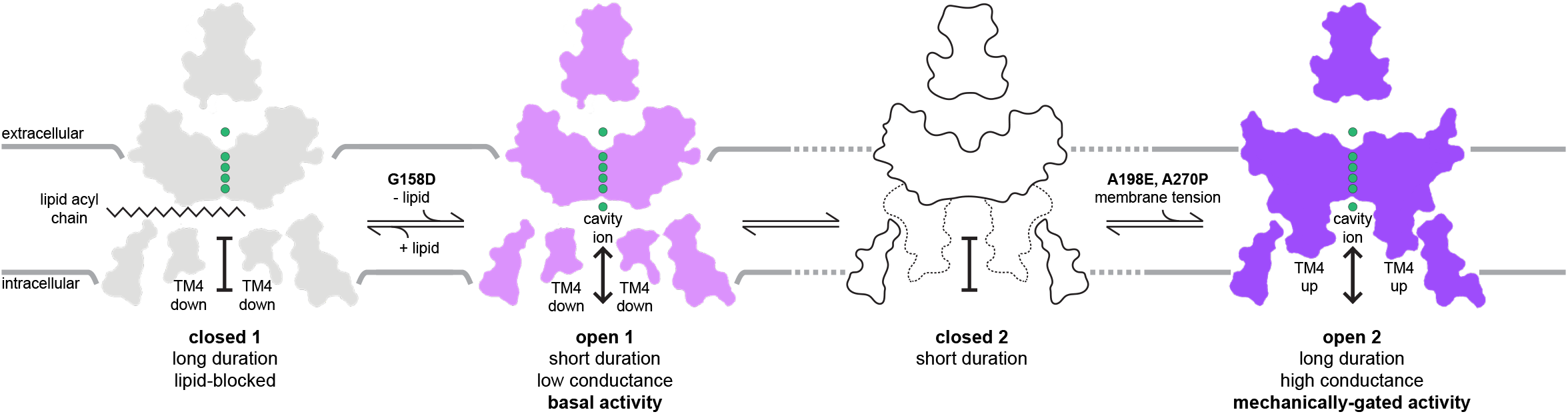
An integrated model for TRAAK gating with distinct basal and mechanically-gated open states. Known structures are mapped to a linear-four state model for TRAAK gating. Basal activity corresponds to a TM4-down, low conductance, short duration open state O1. Mechanically-gated activity corresponds to a TM4-up, high conductance, long duration open state O2. Long duration closures correspond to a TM4-down lipid-blocked state C1. The unknown structure of the short duration closed state C2 is drawn without ions and the position of TM4 is indicated with dashed lines.

TRAAK_WT_ under basal conditions transitions primarily between C1, O1, and C2. Mechanical force favors the TM4 up open state O2 because it involves shape changes (relative to TM4 down) that are energetically favored in the presence of membrane tension, including an increase in cross-sectional area and cylindricity(Brohawn et al., 2014b). Mechanical force disfavors the TM4 down states C1 and O1 because the relatively smaller cross-sectional area and more wedge-shaped structure is disfavored in the presence of membrane tension. The gain of function mutations TRAAK_A198E_ and TRAAK_A270P_ mimic mechanically activated TRAAK_WT_ by promoting a TM4 up conductive conformation. As a consequence, TRAAK_A198E_ and TRAAK_A270P_, like mechanically activated TRAAK_WT_, primarily transition between O2 and C2 states. Gain of function mutant TRAAK_G158D_ mimics basally open TRAAK_WT_, promoting a conductive TM4 down conformation by disfavoring lipid entry and channel block at low membrane tension. As a consequence, TRAAK_G158D_ primarily transitions between O1, C2, and O2 states. Like TRAAK_WT_, mechanical stimulation of TRAAK_G158D_ disfavors O1 to favor transitions between O2 and C2 states.

The short duration closure (C2) does not yet have a defined structural correlate. We observe C2-like closures of similar duration in all TRAAK variants analyzed (Figs. 1L, 2J, 3J, 4K), suggesting they may occur independently of TM4 position or cavity environment. One possibility is that C2 closures represent gating at the selectivity filter. While gating conformational changes at the selectivity filter have not been observed in TRAAK, they have been hypothesized from functional data and visualized in other K2Ps including the mechanosensitive TREK1 in low [K^+^] and pH-gated TASK2 in physiological [K^+^] and low pH (Li et al., 2020; Lolicato et al., 2020). Alternatively, C2 closures could correspond to cavity dewetting that has been proposed to occur based on molecular dynamics simulations (Aryal et al., 2014, 2015) or other, yet to be visualized, conformational rearrangements.

Notably, some evidence suggests that the physical basis for distinct modes of TRAAK activity shown here could be shared among K2P channels. Differences between basal and stimulus-activated openings, including their kinetics and voltage dependence (Schewe et al., 2016), have been reported for other K2Ps, suggesting that physically distinct open states could similarly underlie different modes of channel activity (Bang et al., 2000; Gnatenco et al., 2002; Kang et al., 2005; Maingret et al., 1999; Clausen et al 2020; Sorum et al., 2021). Mutations analogous to TRAAK_G158D_ were found to activate all K2P channels, with the extent of activation correlated with the polarity of the introduced amino acid change (Ben Soussia et al., 2019). Membrane-facing openings have been observed in TM4-down-like structures of TREK2 (Dong et al., 2015), TWIK1 (Miller and Long, 2012), TASK1 (Rödström et al., 2020), and TASK2 (Li et al., 2020) in addition to TRAAK(Brohawn et al., 2012) (Fig. S7D-L). Acyl chains that could block conduction were modeled in TWIK1 and TRAAK structures (Brohawn et al., 2012; Miller and Long, 2012), though their absence in other K2Ps may be due to limited resolution of, or partial occupancy in, structures determined to date. This raises the possibility that lipid-mediated block could be more common among K2Ps than currently appreciated. Like TRAAK, basal activity of other K2Ps could result from periodic unbinding of lipid from channel cavities and ion passage through a transiently unobstructed conduction path, while stimulus gating could involve conformational changes that prevent lipid entry into the channel entirely (Brohawn et al., 2012, 2014b; Dong et al., 2015; Li et al., 2020; Miller and Long, 2012; Rödström et al., 2020).

## Methods

### Electrophysiology

Full length hsTRAAK (Uniprot Q9NYG8-2) was codon optimized for eukaryotic expression, synthesized (Genewiz), and cloned into a modified pGEMHE vector using XhoI and EcoRI restriction sites such that the mRNA transcript encodes hsTRAAK 1-419 with an additional “SNS” at the C-terminus. Point mutants G158D, A198E, and A270P were generated by inverse PCR of the parent construct and verified by Sanger sequencing. cRNA was transcribed from linearized plasmids *in vitro* using T7 polymerase, and 0.1–10 ng cRNA was injected into *Xenopus laevis* oocytes extracted from anaesthetized frogs.

Currents were recorded at 25°C from inside-out patches excised from oocytes 1– 5 days after mRNA injection. Pipette solution contained (in mM) 15 KCl, 135NaCl, 2MgCl_2_, 10 HEPES, pH = 7.4 with NaOH. The solution in the bath contained 150 KCl, 2 MgCl_2_, 10 HEPES, 1 EGTA, pH = 7.1 with KOH. All single channel records were made at a holding voltage of 0 mV and presented in physiological convention.

Currents were recorded using an Axopatch 200B Patch Clamp amplifier at a bandwidth of 1 kHz and digitized with an Axon Digidata 1550B at 50 to 500 kHz. To maintain the integrity of brief TRAAK dwell times, records were not further filtered unless explicitly stated. Baseline correction was performed manually based on the observable closed state for each recording; since there was no solution exchange in our experiments, no associated large drifts in baseline were observed. Portions of TRAAK_WT_ records were assigned as low, mid, or high P_O_ for analysis. Low P_O_ was observed prior to pressure application, while mid and high P_O_ behavior was observed after pressure application. Portions of mutant records during pressure application were analyzed separately from mutant records prior to pressure application. High P_O_ TRAAK_WT_ and mutant records with pressure were made with the highest achievable pressure prior to patch rupture. Single channel open-close transitions were idealized by half-amplitude threshold crossing and dwell time event lists were generated for each baseline corrected record. Dwell time histograms were generated from the dwell time event lists using custom built software (Sorum et al., 2015), while the fits for these histograms were generated in GraphPad Prism version 9.0 using single, double, or triple Gaussian fits.

For unitary current analysis, event lists for TRAAK_WT_ low P_O_, TRAAK_WT_ mid P_O_, TRAAK_G158D_, and TRAAK_G158D_ with pressure were first generated in Clampfit 10.7, applying a 0.8 pA event detection cutoff. Currents were filtered to 10 kHz and were analyzed with a custom script in Python 3.7 using the pyABF module (Harden, 2019). Each point in the filtered record was annotated as closed (C1/C2), open low conductance (O1), or open high conductance (O2) using the events list. To account for the sharp detection cutoff, openings were padded by an additional point on either side of the cutoff (these points corresponded to 100 us of ∼0.5-1 pA current during openings and closings). Open only event lists were then generated by removing all closed points. Open event only lists were binned by 0.2 pA in current and 0.2 ms in time to generate bubble plots relating unitary current and mean open duration for TRAAK_WT_ and TRAAK_G158D_. Open only event lists were also used to generate open event current histograms to distinguish relatively rare low conductance openings from prevalent closed events. Mean unitary currents for TRAAK_WT_ low P_O_ and TRAAK_WT_ mid P_O_ were derived from Gaussian fits to these open event current histograms. Mean unitary currents for TRAAK_WT_ high P_O_ and all mutants were derived from Gaussian fits to square root total event current histograms in order to distinguish conductance states when one state is present at higher frequency.

### TRAAK expression and purification

Human TRAAK UniProt Q9NYG8-2 was cloned for expression in *Pichia pastoris* cells as previously described (Brohawn et al., 2012) with modifications described here. The construct used for purification included an additional 26 amino acid N-terminal sequence compared to Q9NYG8-1 that improved heterologous expression. The final construct is C-terminally truncated by 119 amino acids, incorporates two mutations to remove N-linked glycosylation sites (N104Q/N108Q), and is expressed as a C-terminal PreScission protease-cleavable EGFP-10xHis fusion protein. As a result, there is an additional amino acid sequence of “SNSLEVLFQ” at the C-terminus of the final purified protein after protease cleavage. Recombinant *Pichia* were grown in a 4 liter fermenter and harvested 48-72h after induction with methanol.

30-120g of frozen *Pichia* cells expressing TRAAK were disrupted by milling (Retsch model MM301) 5 times for 3 minutes at 25 Hz. All subsequent purification steps were carried out at 4 °C. Milled cells were resuspended in buffer A (in mM) 50 Tris pH 8.0, 150 KCl, 1 EDTA 0.1 mg/mL DNase1, 1 mg/mL pepstatin, 1 mg/mL leupeptin, 1 mg/mL aprotinin, 10 mg/mL soy trypsin inhibitor, 1 mM benzamidine, 100 µM AEBSF, 1 µM E-64, and 1 mM phenylmethysulfonyl fluoride added immediately before use) at a ratio of 1 g cell pellet per 4 mL lysis buffer and sonicated for 16 minutes with a duty cycle of 15 seconds of sonication per minute. The solution was ultracentrifuged at 150,000 xg for 1 hour at 4 °C. Pellets were transferred to a Dounce homogenizer in buffer B (buffer A + 60 mM DM). Following homogenization, solutions were stirred for 3 hours at 4 °C followed by centrifugation at 33,000g for 45 minutes. Anti-GFP nanobody resin (1 mg purified anti-GFP nanobody conjugated to 1 mL resin) was washed in Buffer B and added to the supernatant at a ratio of 1 mL resin / 15 g *Pichia* cells. The solution was gently stirred for 3 hours. Resin was collected on a column and washed in 15 column volumes (CV) of Buffer C (buffer A +6 mM DM+ 150 mM KCl), followed by 2 CV Buffer D (buffer A +6 mM DM). The resin was resuspended in ∼3 CV of Buffer D with 1 mg purified Precission protease and gently rocked in column overnight. Cleaved TRAAK was eluted in ∼4 CV of Buffer D, concentrated (50 kDa MWCO), and applied to a Superdex 200 SEC column (GE Healthcare) equilibrated in Buffer E (20 mM Tris pH 8.0, 150 mM KCl, 1 mM EDTA, 4 mM DM). Peak fractions were pooled and concentrated to 200-300 µL for incubation with Fab 13E9 and subsequently applied to a Superdex 200 column (GE Healthcare) equilibrated in Buffer E. Fab 13E9 was prepared as described previously(Brohawn et al., 2014b). TRAAK-Fab complexes were pooled and concentrated to 25-33 mg/mL for hanging drop crystallization.

For crystallization in Tl^+^, all purification steps were conduction as with KCl, with two exceptions: KCl is substituted for KNO_3_ for all buffers except Buffer E, where KCl is substituted with TlNO_3_.

### Protein Crystallization

Crystals were grown in drops of 0.125–0.250 µL protein added to an equal volume of reservoir, in hanging drops over a 100 µL reservoir at 4 °C. Reservoir for each mutant in KCl was 50mM Tris pH 8.8, 64-200 mM CaCl_2_, 27–33%(vol/vol) PEG400. Reservoir for each mutant in TlNO_3_ was 50 mM Tris pH 8.8, 64-200 mM Ca(NO3)_2_, 27– 33%(vol/vol)PEG400. Lower Calcium concentrations were observed to correspond to increased nucleation and rapid growth of large, well-ordered crystals. Crystals grew to ∼100 µm x 100 µm x 200 µm in 1–5 weeks. TRAAK A270P crystals rarely achieved sizes larger than 70 µm x 70 µm x 100 µm after 5 weeks.

For cryoprotection, an approximately equal volume of mother liquor supplemented to be 30%(vol/vol) PEG400 was added to one side of the drop and crystals were moved through this solution with a cryoloop before being plunged into liquid nitrogen.

### X-ray Data Collection, Model Building & Refinement

Data were collected at beamline 8.3.1 at the ALS, or 24-IDC & 24-IDE at the APS. Thallium-containing crystals were collected at 12680 eV, and Potassium-containing crystals were collected at 12663 eV. Were rotated 360 degrees, and data were collected in 0.20 degree wedges with a 0.25 second exposure time per wedge. Datasets were processed as individual wedges in XDS (Kabsch, 2010; Kabsch et al., 2010). Some datasets were truncated to eliminate redundant data displaying beam-induced damage prior to scaling with XSCALE, and merging with XDSCONV (Kabsch et al., 2010). Due to the anisotropy of the data, elliptically truncated resolution cutoffs generated by STARANISO (Tickle et al., 2018) were used to increased overall resolution of the maps. Structures were solved by molecular replacement using Phaser (McCoy et al., 2007) with an input model of nonconductive hsTRAAK for G158D (PDB ID 4WFF) and conductive hsTRAAK for A198E and A270P mutants (PDB ID 4WFE). Structure refinement was carried out in Coot (Emsley and Cowtan, 2004) and Refmac5 (Murshudov et al., 2011). Jelly body and automatically generated local NCS constraints were used throughout refinement. For the final round of refinement, three TLS groups per protein chain were incorporated. Molprobity was used to assess model geometry in later stages of refinement (Chen et al., 2010).

Ion occupancy in the conduction axis was determined by calculating either a model phased anomalous difference map for A198E crystals grown in Thallium, or a Polder map for G158D and A270P crystals grown Potassium. In all maps, density in the filter fits well to ions occupying the S1, 2, 3, and 4 positions. Similarly, densities above the filter (S0) are well fit with one or two ions. Density in the cavity below the filter is also fit with an ion. All ion occupancies are set to 1.

The angle and orientation of transmembrane helix 4 in each model was compared using UCSF Chimera (Pettersen et al., 2004). Cross-sectional area calculations were performed with CHARMM (Brooks, B. R. and Brooks, III, C. L. and Mackerell, Jr. et al., 2009; Jo et al., 2007, 2008). The area calculation used a surface calculated with 1.4 Å added to the van der Waals radii of protein atoms, and a probe radius of 3.5 Å to approximate lipid-accessible surface area. A water cylinder with a 6 Å radius was used to fill the cavity of TRAAK channels to exclude its contribution from the calculation. All calculations are made with channels containing symmetric protomers of the B chain, which does not make a contact in the crystal. Symmetric molecules were generated by rotation of the B protomer about the conduction axis, using pore helix and selectivity filter residues 119-133 and 228-242. Pore radii and pore-lining residue hydropathy were calculated with CHAP (Klesse et al., 2019) using a pathway lining residue margin value of 0.55 and a hydrophobicity kernel bandwidth of 0.45.

## Author contributions

S.G.B., R.A.R., and B.S. conceived of the project, analyzed data, and wrote the manuscript. R.A.R. performed all aspects of the biochemistry and structural biology. B.S. performed all aspects of the electrophysiology. R.A.R. and B.S. generated constructs for recording. B.L. cultured the hybridoma for antibody purification. S.G.B. supervised the project and secured funding.

## Data Availability

The X-ray crystallographic coordinates and structure factors for TRAAK_A198E_ in K^+^ (7LJ5), TRAAK_A198E_ in Tl^+^ (7LJA), TRAAK_A270P_ in K^+^ (7LJ4), and TRAAK_G158D_ (7LJB) are available at the Protein Data Bank.

## Acknowledgements

We thank R. MacKinnon for the 13E9 anti-human TRAAK hybridoma line. We thank staff at ALS beamline 8.3.1, especially J. Holton and G. Meigs, at APS beamline 24-IDC/E, especially I. Kourinov, for assistance at the synchrotron. We thank all members of the Brohawn laboratory for discussions. SGB is a New York Stem Cell Foundation-Robertson Neuroscience Investigator. This work was supported by the New York Stem Cell Foundation, NIGMS grant DP2GM123496, a McKnight Foundation Scholar Award, a Klingenstein-Simons Foundation Fellowship Award, a Sloan Research Fellowship, and a Rose Hill Innovator Award to SGB.

---

**Figure S1.**
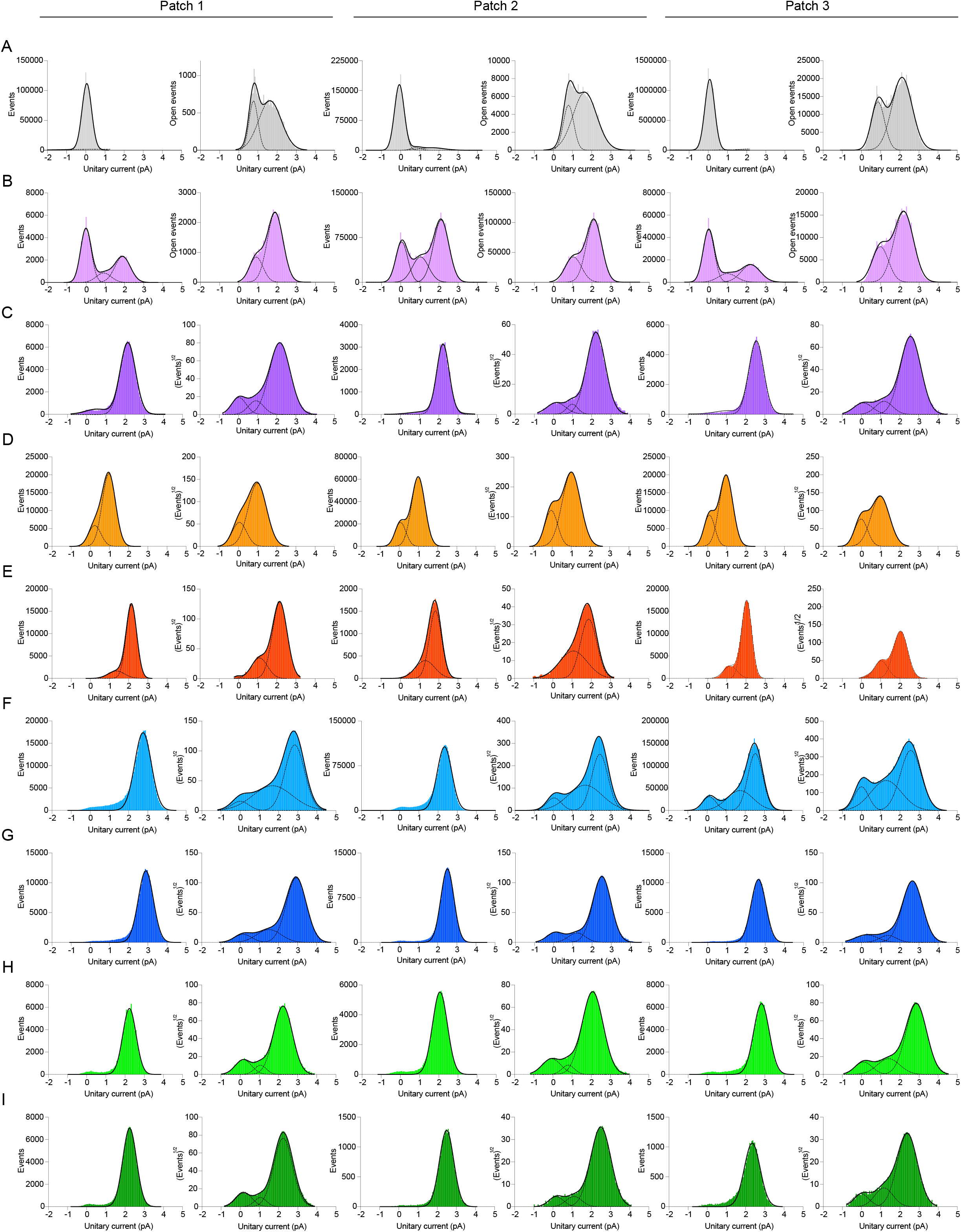
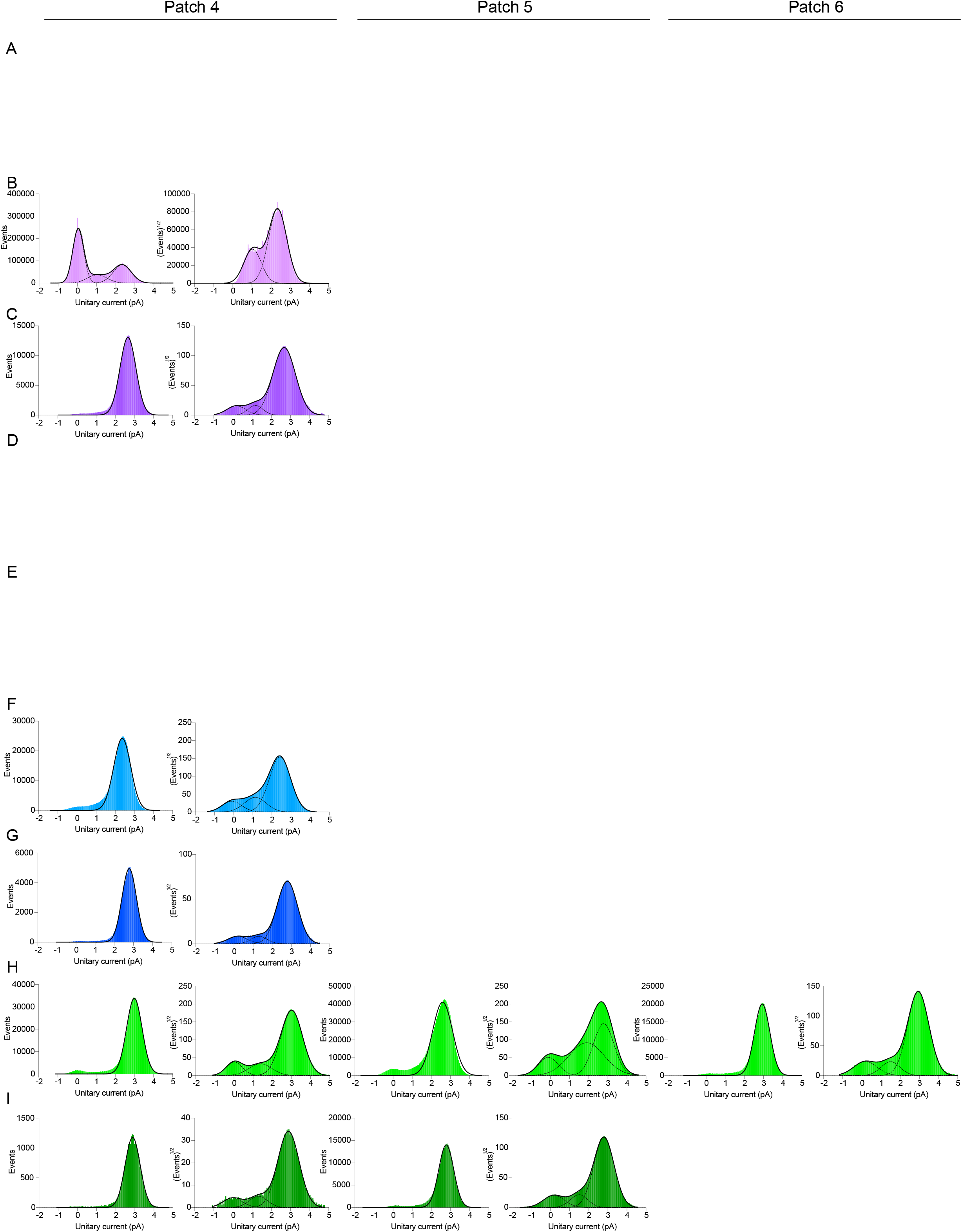
Unitary current histograms. Open event histograms with individual Gaussian fits (dashed lines) and summed fits (solid lines) for each record analyzed. (A) TRAAK_WT_ low P_O_, (B) TRAAK_WT_ mid P_O_, (C) TRAAK_WT_ high P_O_, (D) TRAAK_G158D_, (E) TRAAK_G158D_ with pressure, (F) TRAAK_A198E_, (G) TRAAK_A198E_ with pressure, (H) TRAAK_A270P_, and (I) TRAAK_A270P_ with pressure.

**Figure S2.**
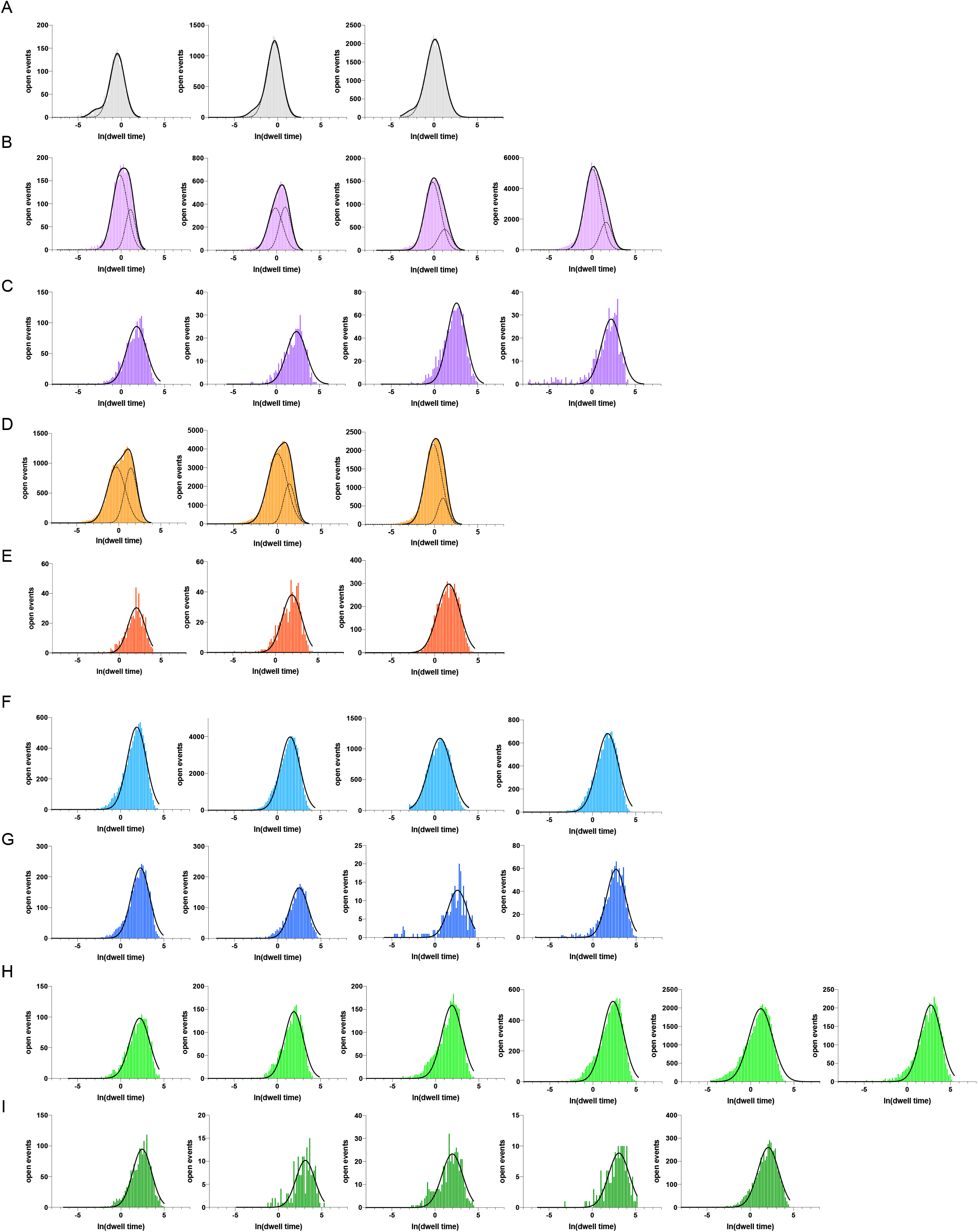
Open event dwell time histograms. Open event histograms with individual Gaussian fits (dashed lines) and summed fits (solid lines) for each record analyzed. (A) TRAAK_WT_ low P_O_, (B) TRAAK_WT_ mid P_O_, (C) TRAAK_WT_ high P_O_, (D) TRAAK_G158D_, (E) TRAAK_G158D_ with pressure, (F) TRAAK_A198E_, (G) TRAAK_A198E_ with pressure, (H) TRAAK_A270P_, and (I) TRAAK_A270P_ with pressure.

**Figure S3.**
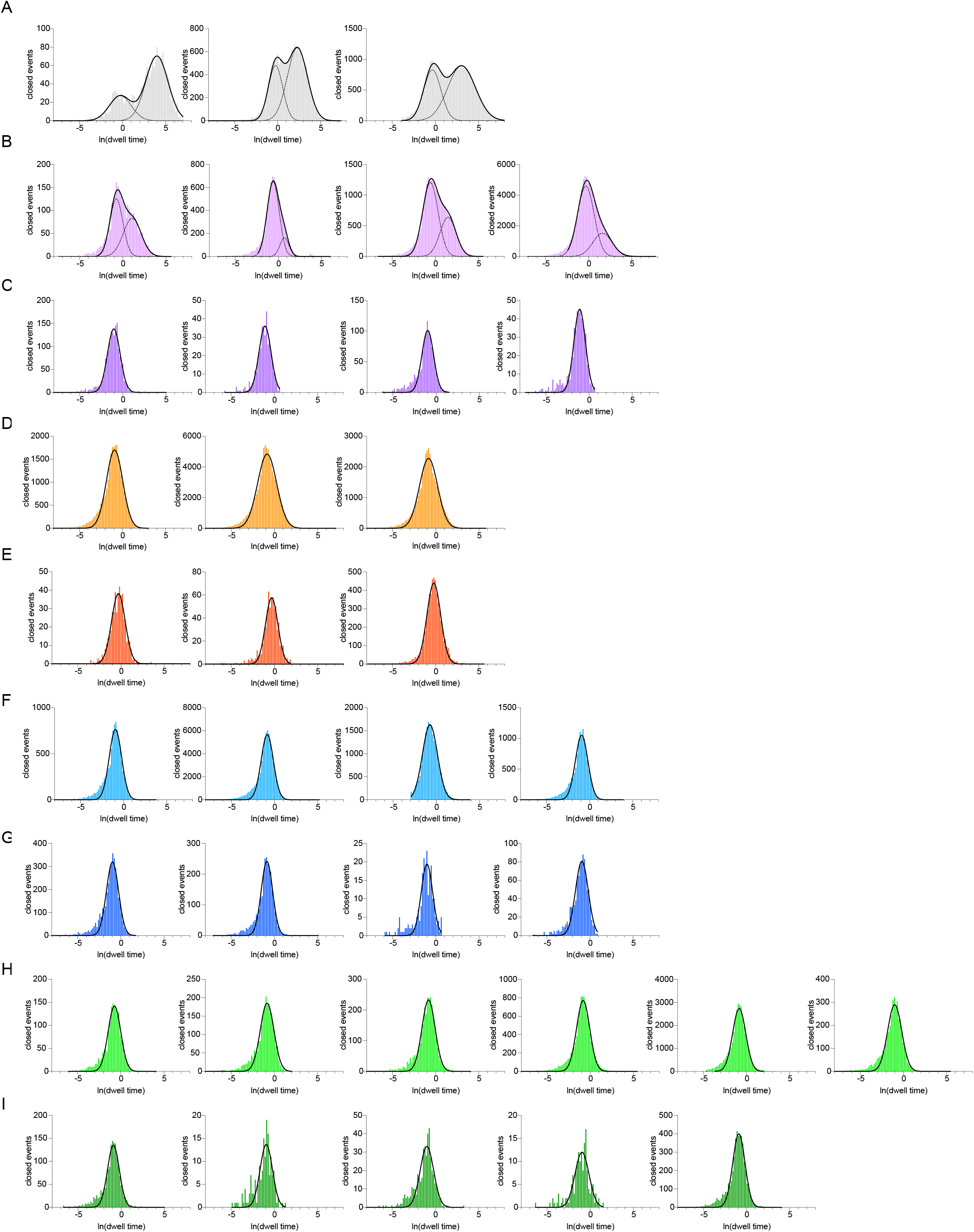
Closed event dwell time histograms. Closed event histograms with individual Gaussian fits (dashed lines) and summed fits (solid lines) for each record analyzed. (A) TRAAK_WT_ low P_O_, (B) TRAAK_WT_ mid P_O_, (C) TRAAK_WT_ high P_O_, (D) TRAAK_G158D_, (E) TRAAK_G158D_ with pressure, (F) TRAAK_A198E_, (G) TRAAK_A198E_ with pressure, (H) TRAAK _A270P_, and (I) TRAAK _A270P_ with pressure.

**Figure S4.**
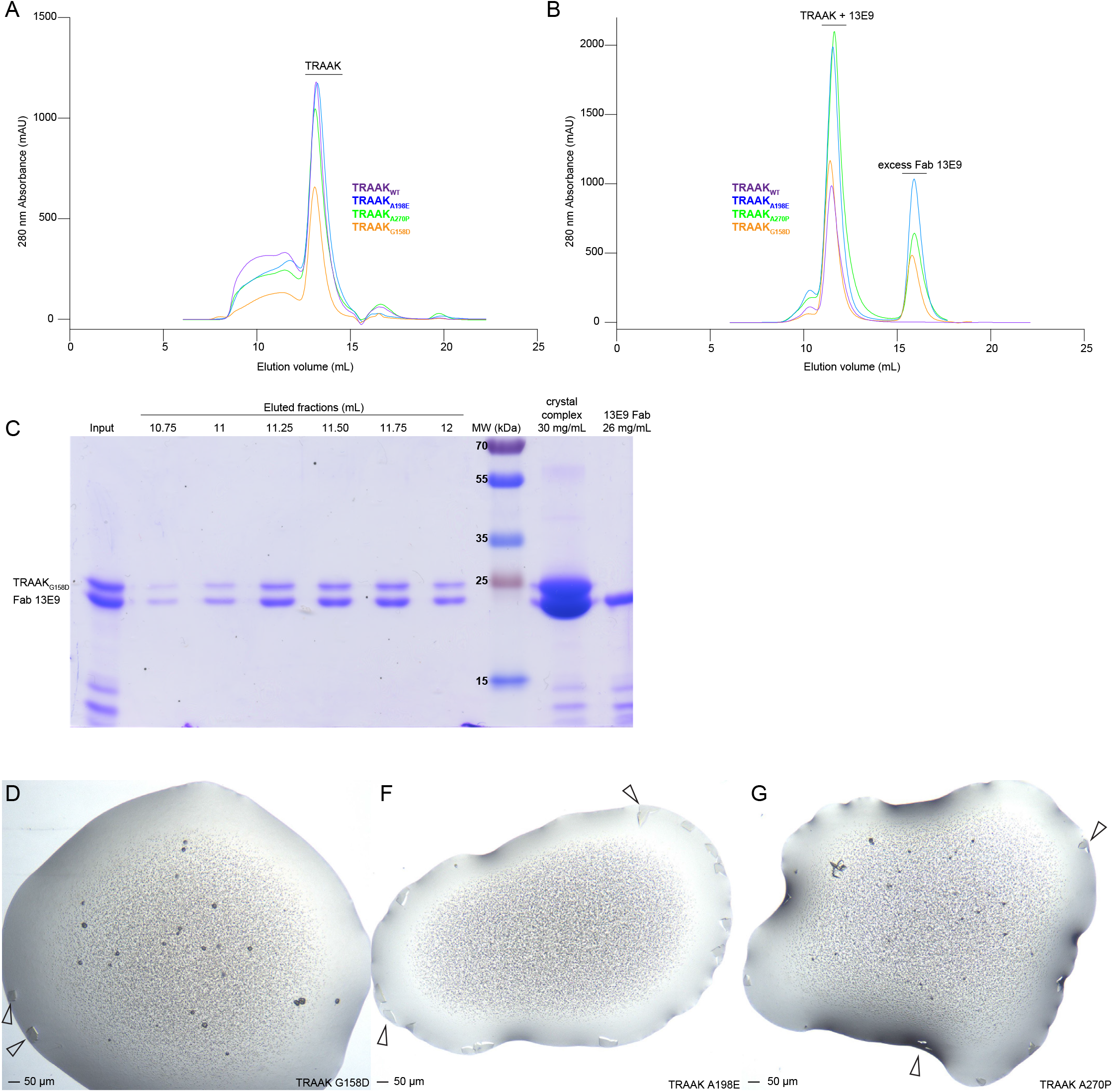
Purification and crystallization of TRAAK_WT_, TRAAK_A198E_, TRAAK_A270P_, and TRAAK_G158D_. (A) Chromatograms from purifications of TRAAK constructs over a Superdex 200 column. (B) Chromatograms purifications of 13E9 Fab – TRAAK construct complexes over a Superdex 200 column. (C) Coomassie-stained SDS-PAGE of fractions from TRAAK_G158D_-13E9 Fab complex purification from (B). (D-G) Representative crystals grown of TRAAK_A198E_, TRAAK_A270P_, and TRAAK_G158D_.

**Figure S5.**
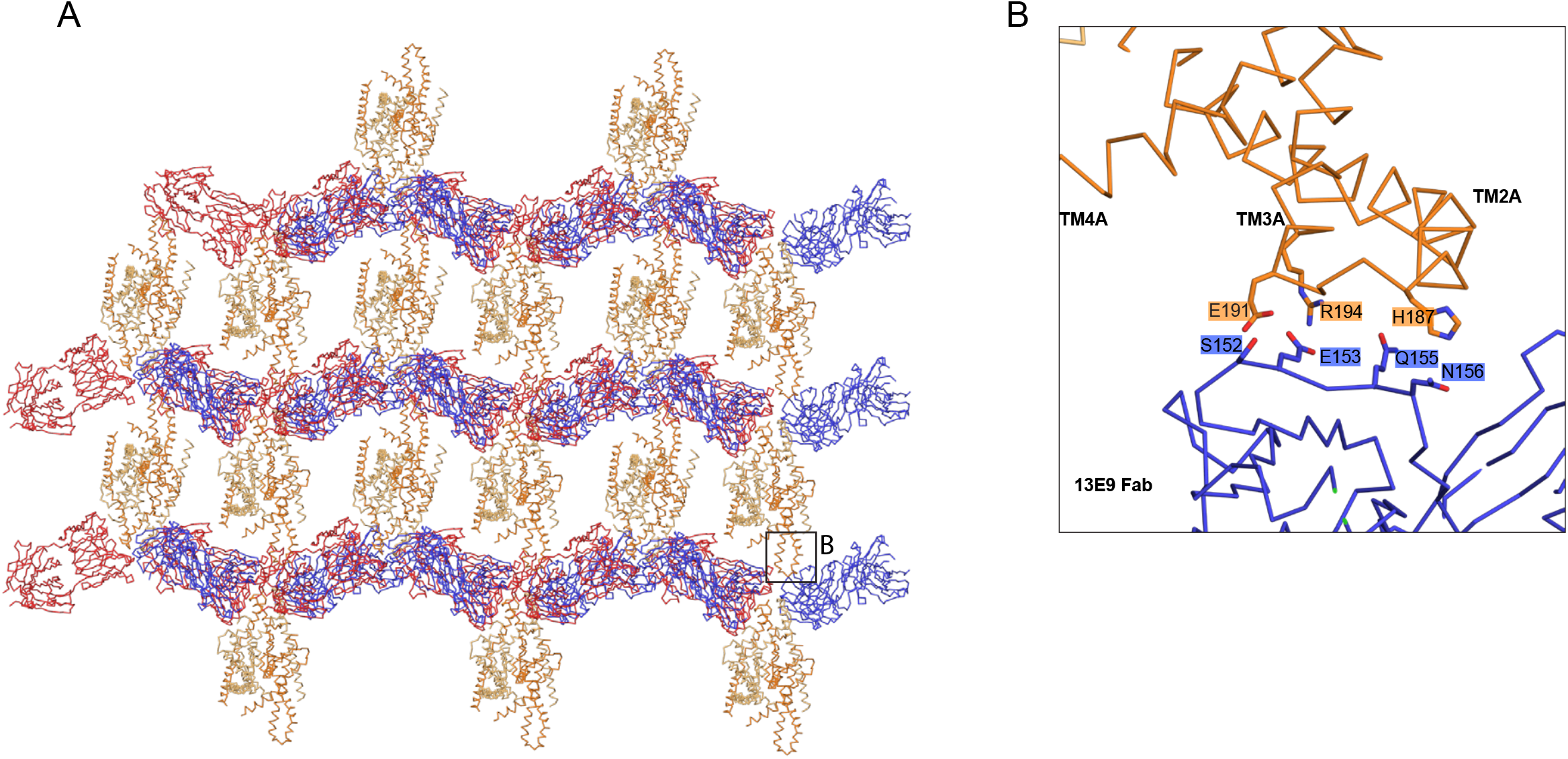
Crystal packing and contacts between TRAAK and monoclonal Fab 13E9. (A) Crystal lattice and (B) zoomed in view of contacts with TRAAK_G158D_ colored orange and Fabs blue and red.

**Figure S6.**
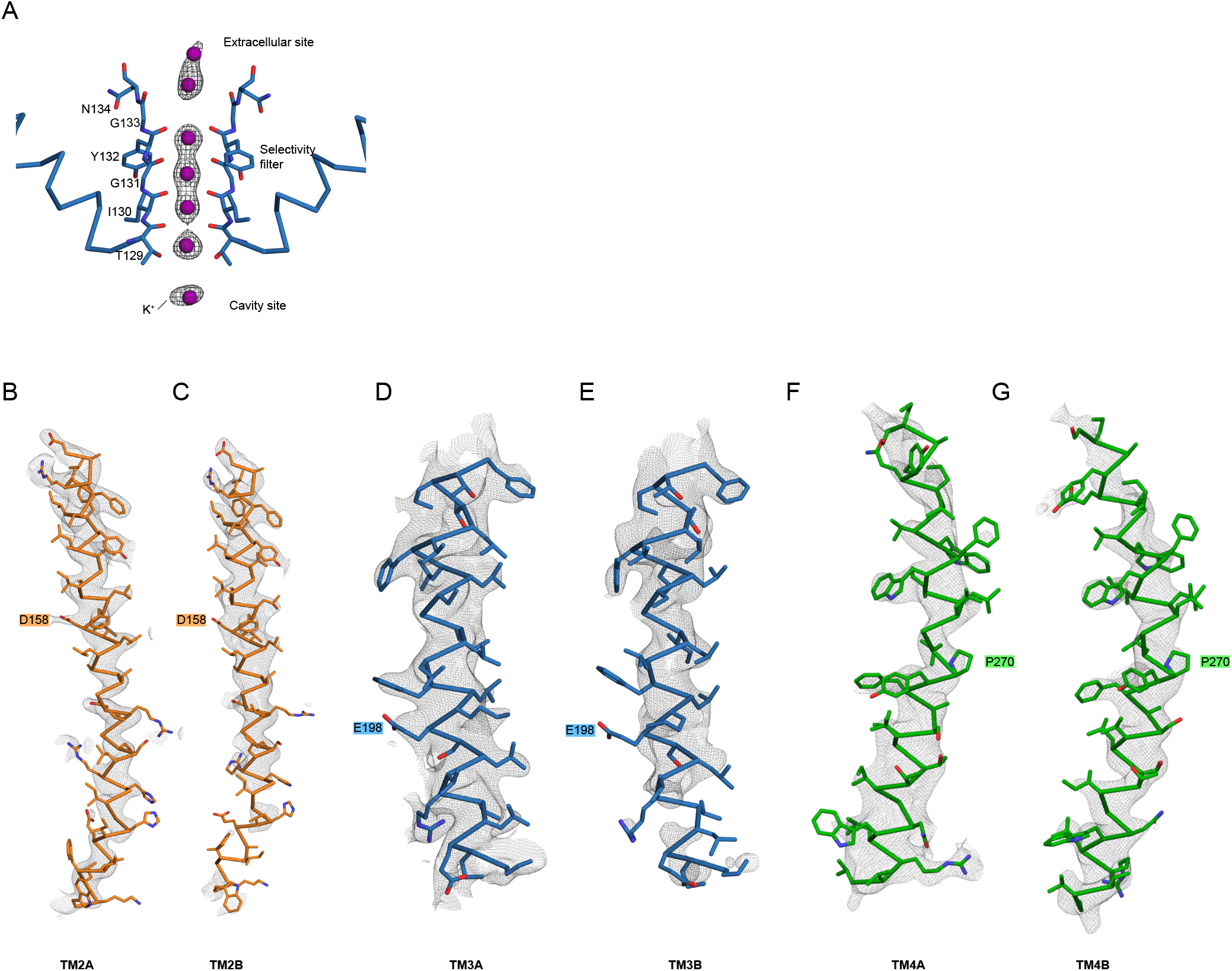
Electron density maps of TRAAK point mutations. (A) Ions in the TRAAK _A198E_-K^+^ structure. Polder omit F _o_-F _c_ density (grey) around K^+^ ions (purple) displayed at 5 and 5.5 σ for selectivity filter and extracellular and cavity ions, respectively. (B-G) TRAAK_G158D_ (orange), TRAAK_A198E_ (blue), and TRAAK_A270P_ (green) models and 2F_o_-F_c_ electron density displayed at 1 σ. Position of mutations are indicated.

**Figure S7.**
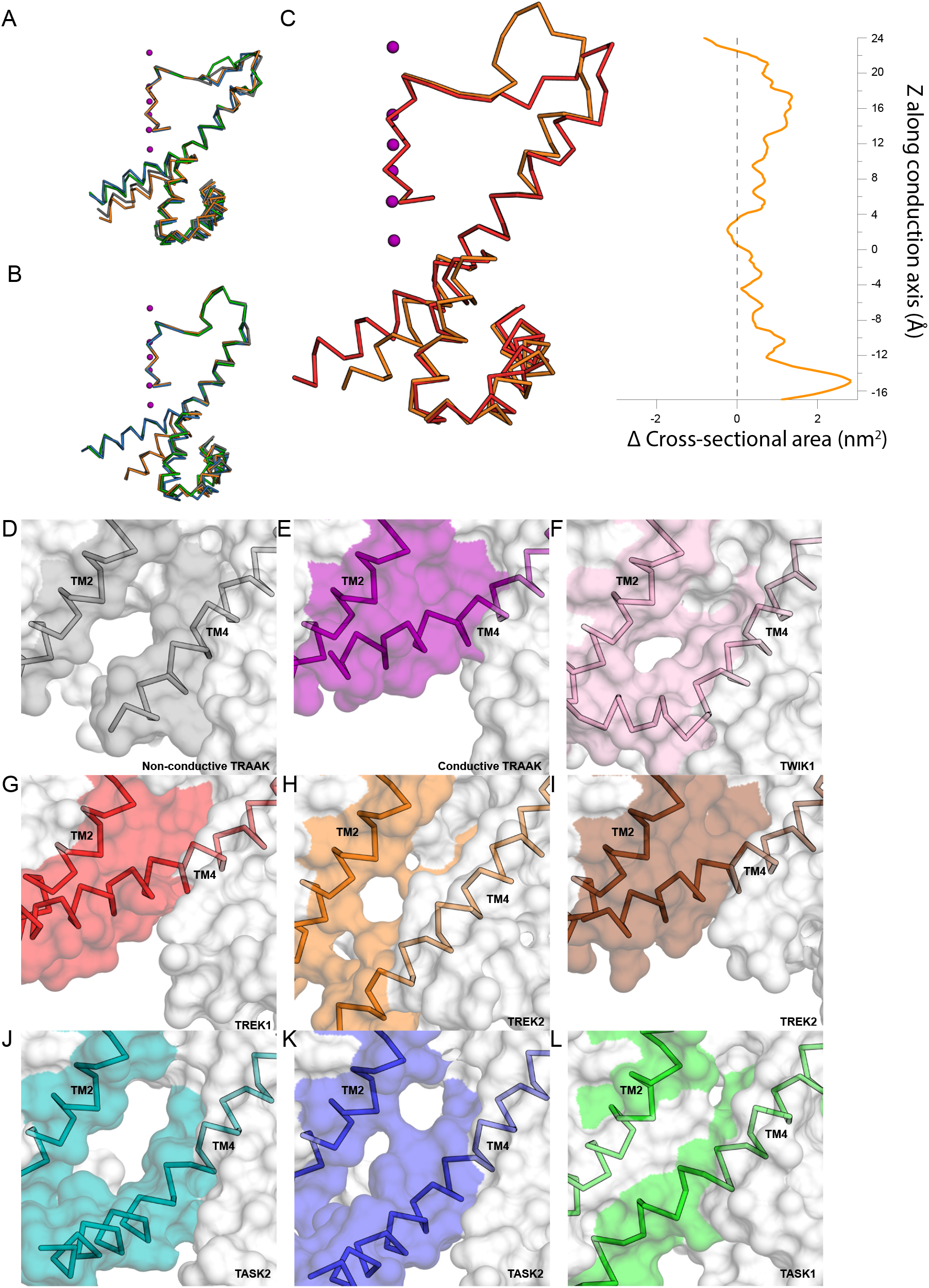
Lateral openings in K2P family ion channels. (A) Overlays of TM4A and (B) TM4B from TRAAK_WT_ (gray), TRAAK_G158D_ (orange), TRAAK_A198E_ (blue), and TRAAK_A270P_ (green) structures crystallized under the same conditions. (C) Change in cross sectional area between symmetric TRAAK_G158D_ TM4-up and TRAAK_G158D_ TM4-down conformations. (D-L) View of the membrane-facing cytoplasmic half of TM4 from K2Ps indicated.

**Table S1.**
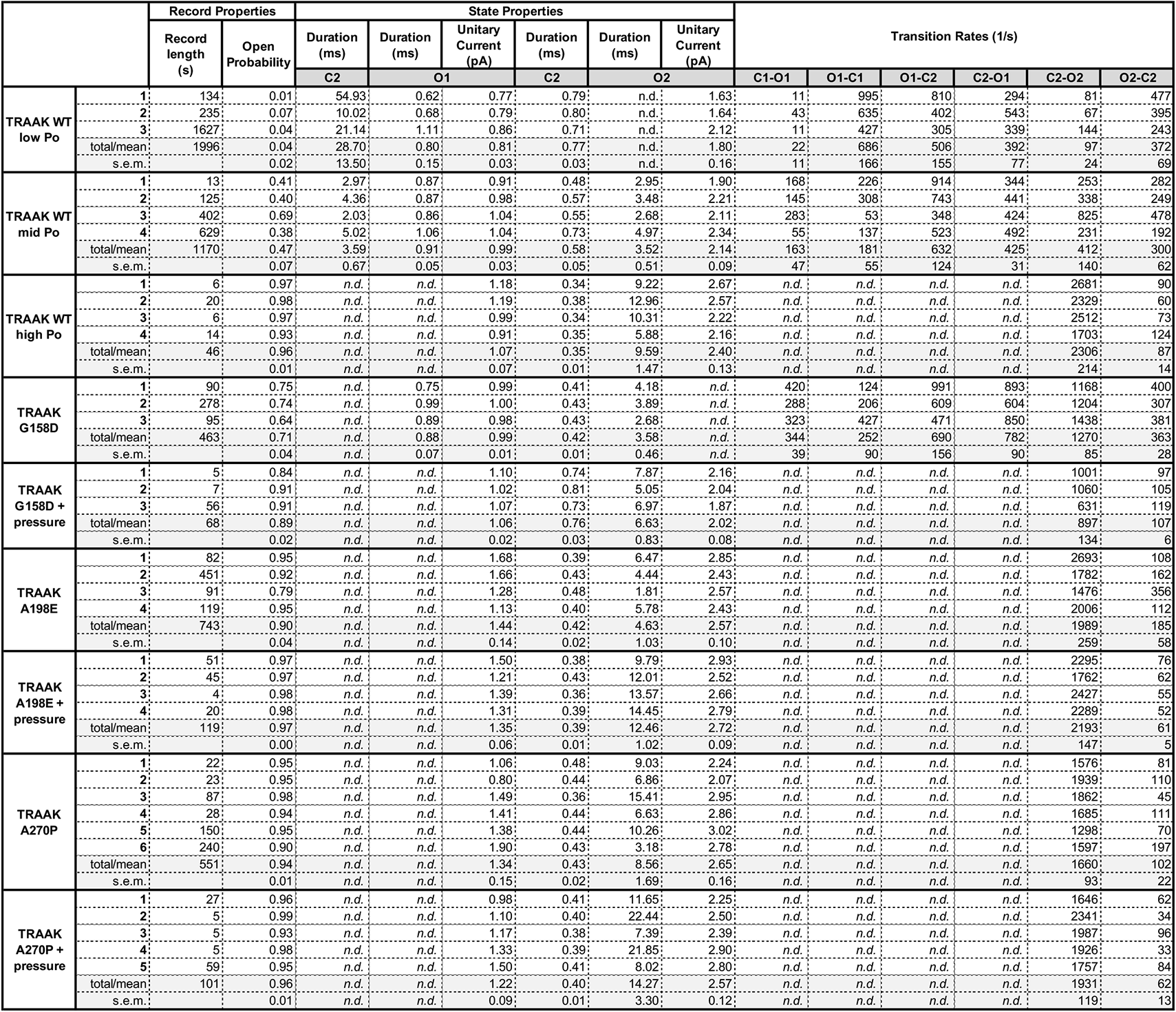
Summary of all single channel recording data.

**Table S2.**
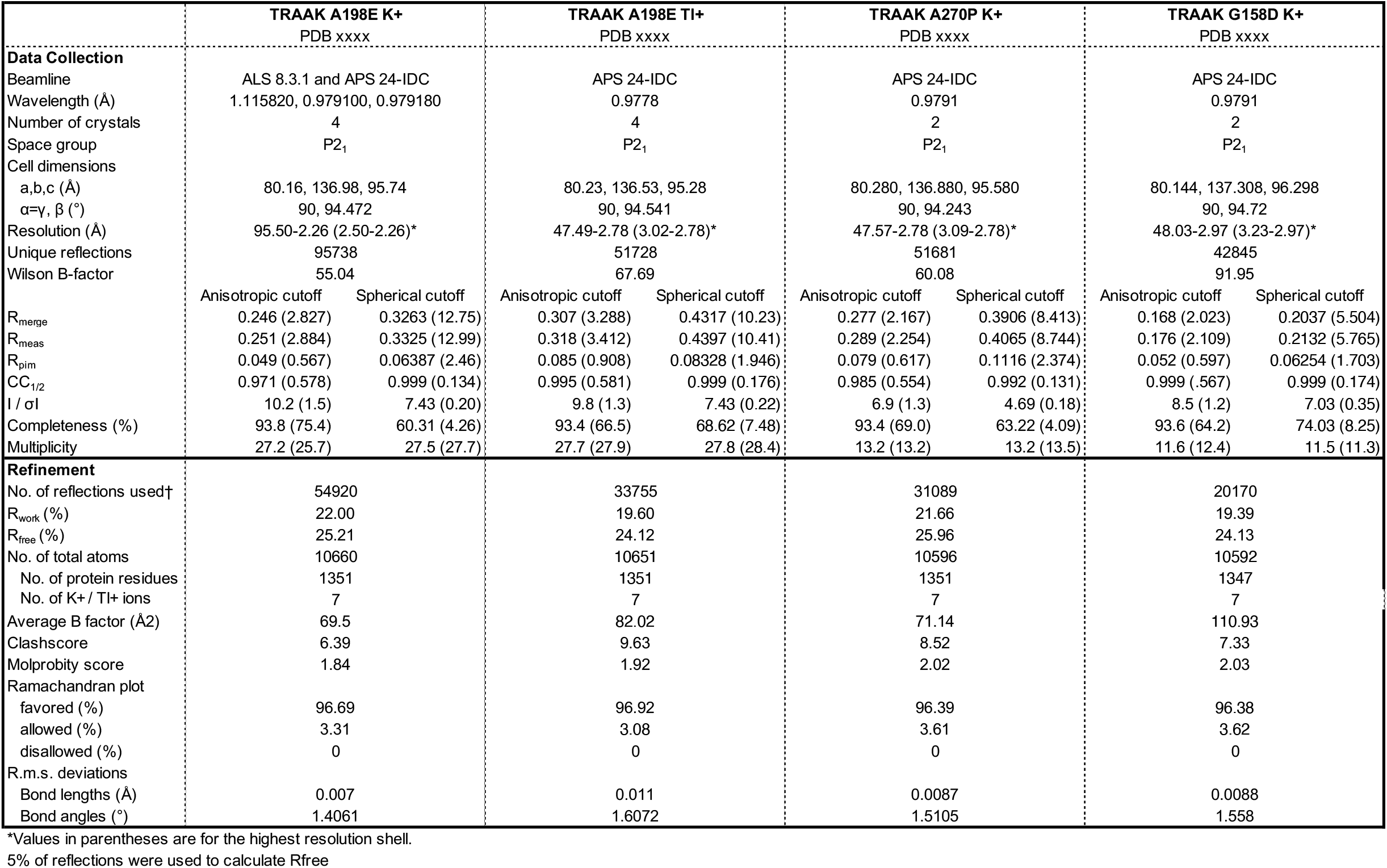
X-ray crystallographic data collection and model refinement statistics.

## References

Aryal, P., Abd-Wahab, F., Bucci, G., Sansom, M.S.P., and Tucker, S.J. (2014). A hydrophobic barrier deep within the inner pore of the TWIK-1 K2P potassium channel. Nat. Commun. 10.1038/ncomms5377.

Aryal, P., Sansom, M.S.P., and Tucker, S.J. (2015). Hydrophobic gating in ion channels. J. Mol. Biol. 10.1016/j.jmb.2014.07.030.

Bang, H., Kim, Y., and Kim, D. (2000). TREK-2, a new member of the mechanosensitive tandem-pore K+ channel family. J. Biol. Chem. 10.1074/jbc.M000445200.

Barel, O., Shalev, S.A., Ofir, R., Cohen, A., Zlotogora, J., Shorer, Z., Mazor, G., Finer, G., Khateeb, S., Zilberberg, N., et al. (2008). Maternally Inherited Birk Barel Mental Retardation Dysmorphism Syndrome Caused by a Mutation in the Genomically Imprinted Potassium Channel KCNK9. Am. J. Hum. Genet. 10.1016/j.ajhg.2008.07.010.

Bauer, C.K., Calligari, P., Radio, F.C., Caputo, V., Dentici, M.L., Falah, N., High, F., Pantaleoni, F., Barresi, S., Ciolfi, A., et al. (2018). Mutations in KCNK4 that Affect Gating Cause a Recognizable Neurodevelopmental Syndrome. Am. J. Hum. Genet. 10.1016/j.ajhg.2018.09.001.

Brohawn, S.G., del Marmol, J., and MacKinnon, R. (2012). Crystal Structure of the Human K2P TRAAK, a Lipid- and Mechano-Sensitive K+ Ion Channel. Science (80-.). 335, 436–441, 10.1126/science.1213808.

Brohawn, S.G., Su, Z., and MacKinnon, R. (2014a). Mechanosensitivity is mediated directly by the lipid membrane in TRAAK and TREK1 K+ channels. Proc. Natl. Acad. Sci. U. S. A. 111, 3614–3619, 10.1073/pnas.1320768111.

Brohawn, S.G., Campbell, E.B., and MacKinnon, R. (2014b). Physical mechanism for gating and mechanosensitivity of the human TRAAK K+ channel. Nature 516, 126–130, 10.1038/nature14013.

Brohawn, S.G., Wang, W., Handler, A., Campbell, E.B., Schwarz, J.R., and MacKinnon, R. (2019). The mechanosensitive ion channel traak is localized to the mammalian node of ranvier. Elife 10.7554/eLife.50403.

Brooks, B. R. and Brooks, III, C. L. and Mackerell, Jr., A.D. and, Nilsson, L. and Petrella, R. J. and Roux, B. and Won, Y. and A.,G. and Bartels, C. and Boresch, S. and Caflisch, A. and Caves, L. and, Cui, Q. and Dinner, A. R. and Feig, M. and Fischer, S. and Gao, J. and, Hodoscek, M. and Im, W. and Kuczera, K. and Lazaridis, T. and Ma, J., and Ovchinnikov, V. and Paci, E. and Pastor, R. W. and Post, C.B. and, Pu, J. Z. and Schaefer, M. and Tidor, B. and Venable, R.M. and, and Woodcock, H. L. and Wu, X. and Yang, W. and York, D. M. and Karplus, M. (2009). CHARMM: The Biomolecular Simulation Program. J. Comput. Chem. 10.1002/jcc.20945

Chatelain, F.C., Bichet, D., Douguet, D., Feliciangeli, S., Bendahhou, S., Reichold, M., Warth, R., Barhanin, J., and Lesage, F. (2012). TWIK1, a unique background channel with variable ion selectivity. Proc. Natl. Acad. Sci. U. S. A. 10.1073/pnas.1201132109.

Chen, V.B., Arendall, W.B., Headd, J.J., Keedy, D.A., Immormino, R.M., Kapral, G.J., Murray, L.W., Richardson, J.S., and Richardson, D.C. (2010). MolProbity: All-atom structure validation for macromolecular crystallography. Acta Crystallogr. Sect. D Biol. Crystallogr. 10.1107/S0907444909042073.

Clausen, M.V., Ulstrup, J., Poulsen, H. and Nissen, P. (2020) The main activatory and tension-sensitive transitions occur within Prozac sensitive down-states of the potassium selective TREK-2 channel. bioRxiv https://doi.org/10.1101/2020.10.22.351205

Dong, Y.Y., Pike, A.C.W.W., Mackenzie, A., McClenaghan, C., Aryal, P., Dong, L., Quigley, A., Grieben, M., Goubin, S., Mukhopadhyay, S., et al. (2015). K2P channel gating mechanisms revealed by structures of TREK-2 and a complex with Prozac. Science (80-.). 347, 1256–1259, 10.1126/science.1261512.

Emsley, P., and Cowtan, K. (2004). Coot: Model-building tools for molecular graphics. Acta Crystallogr. Sect. D Biol. Crystallogr. 10.1107/S0907444904019158.

Enyedi, P., and Czirják, G. (2010). Molecular background of leak K+ currents: two-pore domain potassium channels. Physiol. Rev. 90, 559–605, 10.1152/physrev.00029.2009.

Gnatenco, C., Han, J., Snyder, A.K., and Kim, D. (2002). Functional expression of TREK-2 K+ channel in cultured rat brain astrocytes. Brain Res. 10.1016/S0006-8993(02)02261-8.

González, W., Zúniga, L., Cid, L.P., Arévalo, B., Niemeyer, M.I., and Sepúlveda, F. V. (2013). An extracellular ion pathway plays a central role in the cooperative gating of a K2P K+ channel by extracellular pH. J. Biol. Chem. 10.1074/jbc.M112.445528.

Harden, S. (2019). Analysis of electrophysiological recordings was performed with custom software written for this project using Python 3.7 and the pyABF module.

Heurteaux, C., Lucas, G., Guy, N., El Yacoubi, M., Thümmler, S., Peng, X.D., Noble, F., Blondeau, N., Widmann, C., Borsotto, M., et al. (2006). Deletion of the background potassium channel TREK-1 results in a depression-resistant phenotype. Nat. Neurosci. 10.1038/nn1749.

Jo, S., Kim, T., and Im, W. (2007). Automated builder and database of protein/membrane complexes for molecular dynamics simulations. PLoS One 10.1371/journal.pone.0000880.

Jo, S., Kim, T., Iyer, V.G., and Im, W. (2008). CHARMM-GUI: A web-based graphical user interface for CHARMM. J. Comput. Chem. 10.1002/jcc.20945.

Kabsch, W. (2010). Integration, scaling, space-group assignment and post-refinement. Acta Crystallogr. Sect. D Biol. Crystallogr. 10.1107/S0907444909047374.

Kabsch, W., T., B.A., K., D., A., K.P., K., D., S., M., G., R.R.B., P., E., S., F., K., W., et al. (2010). XDS. Acta Crystallogr. Sect. D Biol. Crystallogr. 10.1107/S0907444909047337.

Kanda, H., Ling, J., Tonomura, S., Noguchi, K., Matalon, S., and Gu, J.G. (2019). TREK-1 and TRAAK Are Principal K+ Channels at the Nodes of Ranvier for Rapid Action Potential Conduction on Mammalian Myelinated Afferent Nerves. Neuron 10.1016/j.neuron.2019.08.042.

Kang, D., Choe, C., and Kim, D. (2005). Thermosensitivity of the two-pore domain K + channels TREK-2 and TRAAK. J Physiol 5641, 103–116, 10.1113/jphysiol.2004.081059.

Klesse, G., Rao, S., Sansom, M.S.P., and Tucker, S.J. (2019). CHAP: A Versatile Tool for the Structural and Functional Annotation of Ion Channel Pores. J. Mol. Biol. 10.1016/j.jmb.2019.06.003.

Lafrenière, R.G., Cader, M.Z., Poulin, J.F., Andres-Enguix, I., Simoneau, M., Gupta, N., Boisvert, K., Lafrenière, F., McLaughlan, S., Dubé, M.P., et al. (2010). A dominant-negative mutation in the TRESK potassium channel is linked to familial migraine with aura. Nat. Med. 10.1038/nm.2216.

Li, B., Rietmeijer, R.A., and Brohawn, S.G. (2020). Structural basis for pH gating of the two-pore domain K+ channel TASK2. Nature 10.1038/s41586-020-2770-2.

Lolicato, M., Riegelhaupt, P.M., Arrigoni, C., Clark, K.A., and Minor, D.L. (2014). Transmembrane helix straightening and buckling underlies activation of mechanosensitive and thermosensitive K_2P_ channels. Neuron 84, 1198–1212, 10.1016/j.neuron.2014.11.017.

Lolicato, M., Natale, A.M., Abderemane-Ali, F., Crottès, D., Capponi, S., Duman, R., Wagner, A., Rosenberg, J.M., Grabe, M., and Minor, D.L. (2020). K2P channel C-type gating involves asymmetric selectivity filter order-disorder transitions. Sci. Adv. 10.1126/sciadv.abc9174.

Ma, L., Roman-Campos, D., Austin, E.D., Eyries, M., Sampson, K.S., Soubrier, F., Germain, M., Treǵouët, D.A., Borczuk, A., Rosenzweig, E.B., et al. (2013). A novel channelopathy in pulmonary arterial hypertension. N. Engl. J. Med. 10.1056/NEJMoa1211097.

Maingret, F., Fosset, M., Lesage, F., Lazdunski, M., and Honoré, E. (1999). TRAAK is a mammalian neuronal mechano-gated K+ channel. J. Biol. Chem. 274, 1381–1387, 10.1074/jbc.274.3.1381.

McCoy, A.J., Grosse-Kunstleve, R.W., Adams, P.D., Winn, M.D., Storoni, L.C., and Read, R.J. (2007). Phaser crystallographic software. J. Appl. Crystallogr. 10.1107/S0021889807021206.

Miller, A.N., and Long, S.B. (2012). Crystal structure of the human two-pore domain potassium channel K2P1. Science 335, 432–436, 10.1126/science.1213274.

Murshudov, G.N., Skubák, P., Lebedev, A.A., Pannu, N.S., Steiner, R.A., Nicholls, R.A., Winn, M.D., Long, F., and Vagin, A.A. (2011). REFMAC5 for the refinement of macromolecular crystal structures. Acta Crystallogr. Sect. D Biol. Crystallogr. 10.1107/S0907444911001314.

Noël, J., Zimmermann, K., Busserolles, J., Deval, E., Alloui, A., Diochot, S., Guy, N., Borsotto, M., Reeh, P., Eschalier, A., et al. (2009). The mechano-activated K+ channels TRAAK and TREK-1 control both warm and cold perception. EMBO J. 10.1038/emboj.2009.57.

Opsahl, L.R., and Webb, W.W. (1994). Lipid-glass adhesion in giga-sealed patch-clamped membranes. Biophys J 66, 75–79, 10.1016/S0006-3495(94)80752-0.

Patel, A.J., Honoré, E., Maingret, F., Lesage, F., Fink, M., Duprat, F., and Lazdunski, M. (1998). A mammalian two pore domain mechano-gated S-like K+ channel. EMBO J. 10.1093/emboj/17.15.4283.

Pettersen, E.F., Goddard, T.D., Huang, C.C., Couch, G.S., Greenblatt, D.M., Meng, E.C., and Ferrin, T.E. (2004). UCSF Chimera - A visualization system for exploratory research and analysis. J. Comput. Chem. 10.1002/jcc.20084.

Renigunta, V., Schlichthörl, G., and Daut, J. (2015). Much more than a leak: Structure and function of K2P-channels. Pflugers Arch. Eur. J. Physiol. 867–894, 10.1007/s00424-015-1703-7.

Rödström, K.E.J., Kiper, A.K., Zhang, W., Rinné, S., Pike, A.C.W., Goldstein, M., Conrad, L.J., Delbeck, M., Hahn, M.G., Meier, H., et al. (2020). A lower X-gate in TASK channels traps inhibitors within the vestibule. Nature 10.1038/s41586-020-2250-8.

Royal, P., Andres-Bilbe, A., Ávalos Prado, P., Verkest, C., Wdziekonski, B., Schaub, S., Baron, A., Lesage, F., Gasull, X., Levitz, J., et al. (2019). Migraine-Associated TRESK Mutations Increase Neuronal Excitability through Alternative Translation Initiation and Inhibition of TREK. Neuron 10.1016/j.neuron.2018.11.039.

Schewe, M., Nematian-Ardestani, E., Sun, H., Musinszki, M., Cordeiro, S., Bucci, G., De Groot, B.L., Tucker, S.J., Rapedius, M., and Baukrowitz, T. (2016). A Non-canonical Voltage-Sensing Mechanism Controls Gating in K2P K+ Channels. Cell 164, 937–949, 10.1016/j.cell.2016.02.002.

Schmidt, C., Wiedmann, F., Voigt, N., Zhou, X.B., Heijman, J., Lang, S., Albert, V., Kallenberger, S., Ruhparwar, A., Szabó, G., et al. (2015). Upregulation of K2P 3.1 K + Current Causes Action Potential Shortening in Patients with Chronic Atrial Fibrillation. Circulation 10.1161/CIRCULATIONAHA.114.012657.

Sorum, B., Czégé, D., and Csanády, L. (2015). Timing of CFTR Pore Opening and Structure of Its Transition State. Cell 10.1016/j.cell.2015.09.052.

Sorum, B., Rietmeijer, R.A., Gopakumar, K., Adesnik, H., and Brohawn, S.G. (2021). Ultrasound activates mechanosensitive TRAAK K+ channels directly through the lipid membrane. Proceedings of the National Acadamy of Sciences https://doi.org/10.1073/pnas.2006980118.

Ben Soussia, I., El Mouridi, S., Kang, D., Leclercq-Blondel, A., Khoubza, L., Tardy, P., Zariohi, N., Gendrel, M., Lesage, F., Kim, E.J., et al. (2019). Mutation of a single residue promotes gating of vertebrate and invertebrate two-pore domain potassium channels. Nat. Commun. 10.1038/s41467-019-08710-3.

Tickle, I.J., Flensburg, C., Keller, P., Paciorek, W., Sharff, A., Vonrhein, C., and Bricogne, G. (2018). STARANISO. Cambridge, United Kingdom Glob. Phasing Ltd.

Vierra, N.C., Dadi, P.K., Jeong, I., Dickerson, M., Powell, D.R., and Jacobson, D.A. (2015). Type 2 diabetes-associated K+ channel TALK-1 modulates β-cell electrical excitability, second-phase insulin secretion, and glucose homeostasis. Diabetes 10.2337/db15-0280.

Xian Tao Li, Dyachenko, V., Zuzarte, M., Putzke, C., Preisig-Müller, R., Isenberg, G., and Daut, J. (2006). The stretch-activated potassium channel TREK-1 in rat cardiac ventricular muscle. Cardiovasc. Res. 10.1016/j.cardiores.2005.08.018.

